# TIR immune signaling is blocked by phosphorylation to maintain growth in plants and animals

**DOI:** 10.1101/2025.04.09.647920

**Authors:** Jun Li, Sisi Chen, Bo Yu, Qingzhong Li, Ruijia Liu, Zaiqing Wang, Li Wan, Yang Zhao

**Affiliations:** Shanghai Center for Plant Stress Biology, CAS Center for Excellence in Molecular Plant Sciences, Chinese Academy of Sciences, Shanghai 200032, China; Key Laboratory of Plant Carbon Capture, Chinese Academy of Sciences, Shanghai 200032, China; National Key Laboratory of Plant Molecular Genetics, CAS Center for Excellence in Molecular Plant Sciences, Institute of Plant Physiology and Ecology, Chinese Academy of Sciences, Shanghai 200032, China; University of Chinese Academy of Sciences, Beijing 100049, China

**Keywords:** TIR domain proteins, attenuation of TIR signaling, phosphorylation, TIR NADase activity, plant growth

## Abstract

Toll/interleukin-1 receptor (TIR) domain proteins are immune signaling components and function as NAD^+^-cleaving enzymes to activate defense responses. Activation of TIRs represses growth and drives cell death in plants and promotes axon degeneration in animals, but how plant TIRs are repressed remains unclear. Here, we show that TIR NADase activity requires a conserved serine residue spatially close to the catalytic glutamate. The plant Ca^2+^-dependent protein kinases (CPKs), the mammalian Ca^2+^/calmodulin-dependent protein kinase II delta (CAMK2D) and TANK binding kinase 1 (TBK1) phosphorylate TIR domains at this conserved serine, which blocks TIR NADase activities and functions and thus maintains growth in plants and suppresses SARM1 TIR signaling in animals, respectively. Our findings define a fundamental molecular mechanism by which phosphorylation at a conserved serine residue blocks TIR signaling to balance growth and defense trade-offs.

## Main

Toll/interleukin-1 receptor (TIR) domain proteins are evolutionarily conserved immune signaling components in prokaryotes and eukaryotes that cleave NAD^+^ (nicotinamide adenine dinucleotide in its oxidized form) and drive cell death^1-4^. TIR domains from plants and bacteria use NAD^+^ as a substrate to generate small signaling molecules that bind and activate downstream proteins^5-8^. In animals, TIR enzymatic activity of the Sterile alpha and TIR motif containing 1 (SARM1) is required for axon degeneration that is associated with a wide array of neurodegenerative diseases, such as amyotrophic lateral sclerosis (ALS)^3,4,9,10^. In plants, TIR domains confer immunity and antagonize hyperosmotic stress resistance and plant growth, although its function in osmotic stress is still unclear^11-13^. Thus, TIR enzymatic activities require delicate and dynamic regulation to maintain plant growth and neuronal function. In animals, nicotinamide mononucleotide adenylyltransferase 2 (NMNAT2) and NAD^+^ repress SARM1 activation and functions^14-16^; however, negative regulators that directly repress plant TIR enzymatic activity remain to be identified.

As a prominent secondary messenger, Ca^2+^ signals mediate diverse abiotic and biotic stress responses, and disruption of Ca^2+^ signals is often associated with altered immune phenotypes^11,17,18^. The phospholipid-binding copine protein BONZAI1 (BON1) modulates cell surface signaling and Ca^2+^ signals, and represses TIR immune signaling mediated by a typical TIR-type intracellular nucleotide-binding leucine-rich repeat (NLR) protein SUPPRESSOR OF NPR1-1, CONSTITUTIVE 1 (SNC1) and potentially several other TIR-NLRs under osmotic stress conditions^11,19-22^. Here we further analyzed how BON1 represses TIR signaling and identified regulators that directly block TIR enzymatic activities. The osmotic stress-activated CPK3 phosphorylates plant TIRs at a conserved serine residue spatially close to the catalytic glutamate, which blocks their NADase activities and immune function. Moreover, the human TBK1 and CAMK2D protein kinases phosphorylate SARM1 TIR and repress its enzymatic activity and cell death phenotype. This study thus uncovers a mechanism underlying the attenuation of TIR signaling and provides a plausible explanation for the balance and switch between defense and growth that is conserved in plants and animals.

### CPK3 represses SNC1-mediated NLR signaling to restore plant growth

Activation of TIR-NLRs severely represses plant growth under osmotic stress^11-13^. The autoimmune *bon1-7* mutant was identified from a mutant library generated in the *AEQsig6* reporter strain in *Arabidopsis thaliana* that has enhanced Ca^2+^ luminescence, and the *bon1-7* mutant is defective in Ca^2+^ signals and plant growth under osmotic stress^11^. To search for additional regulators involved in BON1-mediated repression of TIR signaling, we screened for suppressors of the *bon1-7* mutant based on plant growth under osmotic stress. Whole-genome resequencing revealed six mutant alleles of *SNC1* named *insensitive to osmotic stress* (*ios*) (Extended Data Fig. S1a). These mutations were distributed on the NB and LRR, and the linker between the two domains (NL linker), probably affecting SNC1 oligomerization, autoinhibition, or activation. Although the *snc1* T-DNA knockout mutant allele partially suppressed the *bon1* mutant^11^, the dominant-negative *ios22* (*snc1-16*) and *ios26* (*snc1-17*) mutants completely suppressed the *bon1* mutant phenotypes (Extended Data Fig. S1b-k). Moreover, the overexpression of *SNC1-G1343D* (corresponding to *snc1-16* mutation) or *SNC1-V518I* (corresponding to *snc1-17* mutation) repressed the growth retardation and autoimmune phenotypes of the *bon1* mutants under osmotic stress or in soil (Fig. 1a, and Extended Data Fig. S1b-k). These data suggest that repression of the typical TIR-NLR protein SNC1 by BON1 is critical for plant growth maintenance. Besides, an autoimmune gain-of-function mutant, *snc1-1*, exhibited severe shoot growth defects under hyperosmotic stress (Extended Data Fig. S1l), further supporting growth repression by SNC1 activation under osmotic stress. Although the plant growth defect of the *bon1-7* mutant was recovered in the *ios102* (*snc1-18*) mutant seedlings, the Ca^2+^ signaling phenotype of the *bon1-7* mutant did not recover, however (Extended Data Fig. S1m), suggesting that *SNC1* may function downstream of Ca^2+^ signals under osmotic stress.

**Fig. 1.**
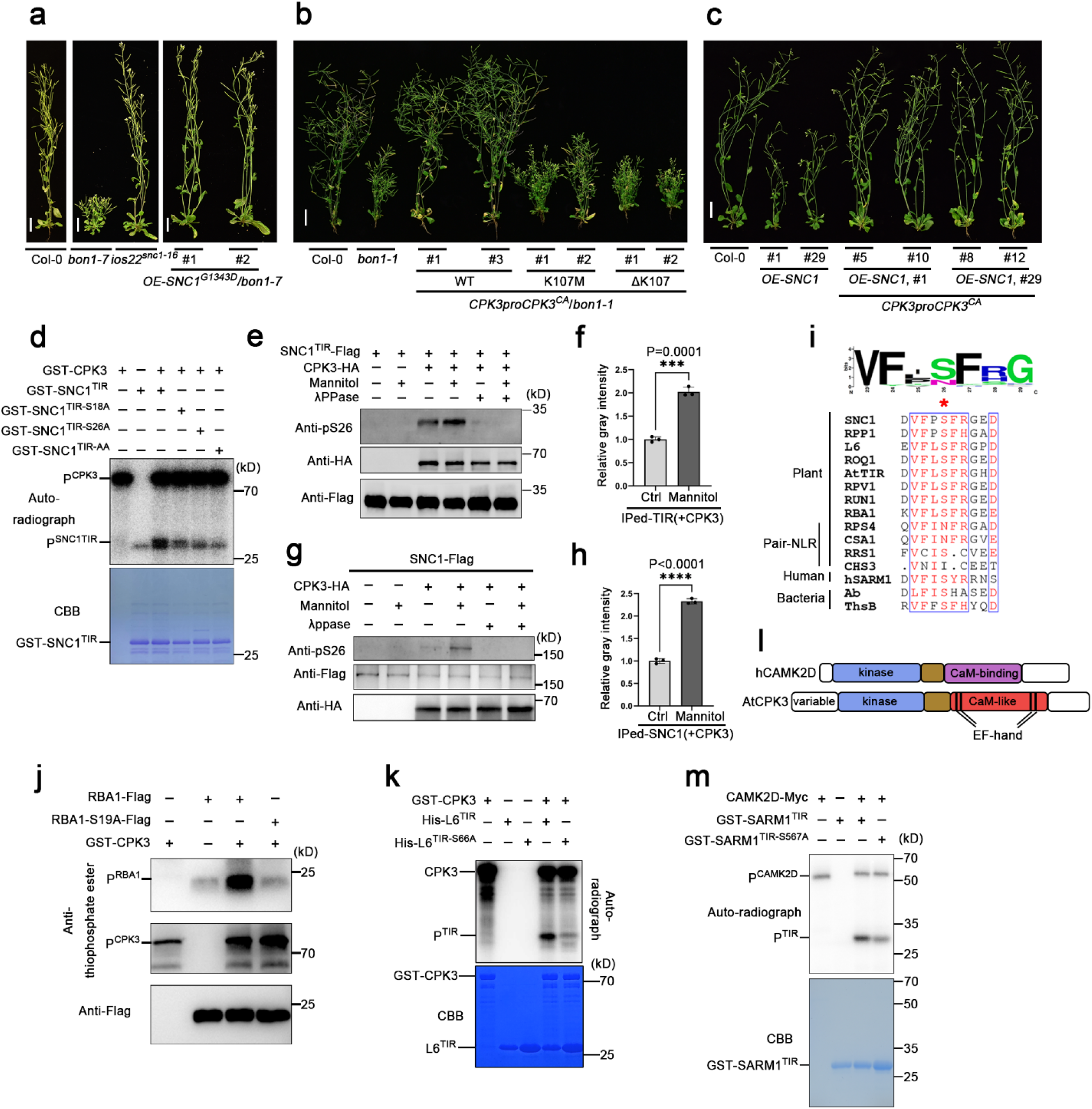
CPK3 phosphorylates a conserved serine in the TIR domain and represses TIR signaling to restore plant growth. **a**, Morphology of 6-week-old soil-grown Col-0 (wild-type), *bon1-7* and *ios22* (*snc1-16*) mutants, and *OE-SNC1*^*G1343D*^ (corresponding to *snc1-16* mutation) transgenic plants in a *bon1-7* mutant background. Scale bars, 2 cm. **b**, Morphology of 6-week-old soil-grown Col-0, *bon1-1* mutant, and *CPK3pro::CPK3*^*CA*^ transgenic plants in a *bon1-1* mutant background, without or with the kinase-dead mutations, including K107M and ΔK107. Scale bars, 2 cm. **c**, Morphology of 5-week-old soil-grown plants of Col-0, and *OE-SNC1-4Myc* transgenic plants with or without *CPK3pro::CPK3*^*CA*^ in Col-0 background. Transgenic plants co-expressing *SNC1-4Myc* and *CPK3*^*CA*^ were generated by crossing two independent transgenic lines of *OE-SNC1-4Myc* and *CPK3pro::CPK3*^*CA*^. Scale bars, 2 cm. **d**, Phosphorylation of wild-type and mutated SNC1^TIR^ by CPK3. Autoradiography (top) and Coomassie staining (bottom) exhibit phosphorylation and the loading of recombinant GST-CPK3, GST-SNC1^TIR^, GST-SNC1^TIR-S18A^, GST-SNC1^TIR-S26A^, and GST-SNC1^TIR-AA (S18A/S26A)^. **e**-**h**, Phosphorylation of Ser26 of SNC1^TIR^-Flag (**e** and **f**) and full-length SNC1-Flag (**g** and **h**) by CPK3-HA *in planta* with or without mannitol treatment. Flag-tagged SNC1^TIR^ or SNC1 was immunoprecipitated from *Nicotiana benthamiana* (*Nb*) leaves co-expressed *SNC1*^*TIR*^*-Flag* or *SNC1-Flag* and *CPK3-HA*, treated with 0 M or 1 M mannitol for 2 hours. Anti-phospho-Ser26-SNC1^TIR^ antibody (anti-pS26) was used to detect the phosphorylation of SNC1^TIR^-Flag (**e**) and SNC1-Flag (**g**). Anti-Flag and anti-HA were used to detect the loading of SNC1^TIR^-Flag or SNC1-Flag, and CPK3-HA, respectively. The relative gray intensity of the bands detected by anti-pS26 was quantified from three independent results (**f** and **h**). **i**, Web-Logo analysis of 141 TIR domains of *Arabidopsis* and multiple sequence alignment of various TIR domains shows a conserved serine phosphosite across kingdoms. **j**, Phosphorylation of wild-type and mutated RBA1 by CPK3. *In vitro* kinase assays were performed using the ATP analog ATPγS. After the PNBM alkylation reaction, the thiophosphate ester groups on the substrate were detected by the anti-thiophosphate ester antibody (top). Anti-Flag was used to detect the loading of RBA1-Flag and RBA1-S19A-Flag (bottom). **k**, Phosphorylation of wild-type and mutated L6^TIR^ by CPK3. Autoradiography (top) and Coomassie staining (bottom) exhibit phosphorylation and the loading of recombinant GST-CPK3, His-L6^TIR^, and His-L6^TIR-S66A^. **l**, Domain comparisons of human CAMK2D and AtCPK3. **m**, Phosphorylation of recombinant wild-type and mutated hSARM1^TIR^ by hCAMK2D. Myc-CAMK2D was immunoprecipitated from HEK293T. Autoradiography (top) and Coomassie staining (bottom) exhibit phosphorylation and the loading of hCAMK2D-Myc, GST-hSARM1^TIR^, and GST-hSARM1^TIR-S567A^. Error bars represent means ± SD (n = 3 experiments) for (**f** and **h**). The statistical analysis was performed using Student's t test. All experiments except (**i** and **l**) were repeated at least three times with similar results.

Given that BON1 controls Ca^2+^ signals and osmotic stress signaling^11,23^, we hypothesized that Ca^2+^ and osmotic stress signaling may modulate SNC1-mediated immune signaling and plant growth. The subgroups I and II CPKs are activated by Ca^2+^ signals upon abiotic stresses and pathogen infections^24-26^, and some of them, such as CPK3, CPK4, CPK6, and CPK11, regulate stomatal closure and seed dormancy that are critical for drought resistance^24,27-30^. Using immunoprecipitation (IP) followed by mass spectrometry (MS) from osmotic stress-treated transgenic seedlings overexpressing *SNC1-Myc*, we were able to identify peptides present in twelve CPKs, among which CPK3 has the most abundant peptides (Extended Data Fig. S1n and Supplementary Table 1). Since *CPK3* is responsive to osmotic stress based on phosphoproteomics data (Extended Data Fig. S1o), is ubiquitously and actively expressed in vegetative tissues and interacts with SNC1 (Extended Data Fig. S1n, p), and is phosphorylated under hyperosmotic stress (Extended Data Fig. S1q-s), we selected CPK3 for further analysis. CPK activities are repressed by their autoinhibitory junction domains, and deletion of the carboxy-terminal Ca^2+^-binding and junction domains generates the constitutively active form of CPKs^25^.

The expression of the native promoter-driven constitutively active form of *CPK3* (*CPK3-CA*) repressed both the hyperactivated TIR-NLR signaling and retarded shoot growth phenotypes in the *bon1-1* mutant under osmotic stress or in soil (Fig. 1b, and Extended Data Fig. S1b, f and t-v). *bon1-1* was chosen because of its less complex background than *bon1-7*^11^. In contrast, the kinase-dead mutations of *CPK3-CA*, including K107M and ΔK107, failed to repress the retarded shoot growth phenotypes in the *bon1-1* mutant (Fig. 1b, and Extended Data Fig. S1x, y). Moreover, the expression of the native promoter-driven *CPK3-CA* repressed the retarded plant growth phenotypes of the *SNC1* overexpression transgenic lines (Fig. 1c and Extended Data Fig. S1z). These data suggest that CPK3 can suppress SNC1 and SNC1-related TIR-NLR signaling, which depends on its kinase activity.

### Phosphorylation of a conserved serine in TIR domains in plants and animals

We next investigated how CPKs are involved in repressing TIR-NLRs and focused on the TIR signaling domain because it is critical for TIR-NLR signaling. First, we tested the potential interactions between SNC1^TIR^ and CPK3. The transient co-expression of *CPK3* and *SNC1*^*TIR*^ generated strong co-immunoprecipitation and reconstituted split luciferase (LUC) signals in *Nicotiana benthamiana* (*Nb*) leaves (Extended Data Fig. S2a, b), which does not require kinase activity (Extended Data Fig. S2c). Second, we investigated the potential regulation of SNC1^TIR^ by CPK3. According to kinase assays and mass spectrometric analyses using purified proteins, we found that recombinant CPK3 phosphorylates the truncated SNC1^TIR^ protein at Ser18 and Ser26 (Fig. 1d, and Extended Data Fig. S2d-g) and the CPK3-mediated phosphorylation of SNC1^TIR^ and full-length SNC1 at Ser26 is enhanced by osmotic stress in *Nb* leaves (Fig. 1e-h, and Extended Data Fig. S2h-j); the Ser26 residue is highly conserved in TIR domain–containing proteins across kingdoms (Fig. 1i), except two plant TIR-NLR pairs examined^31-33^. Mutation of Ser18 and Ser26 to non-phosphorylatable alanines (A) abolished the phosphorylation of SNC1^TIR^ by CPK3 (Fig. 1d and Extended Data Fig. S2f). The phosphorylation of Ser26 was also detected from the IP-MS assays from *Nb* leaves transiently expressed SNC1^TIR^ and CPK3^CA^ (Extended Data Fig. S2k). The phosphorylation of recombinant SNC1^TIR^ was further verified using immunoprecipitated CPK3 from *CPK3pro::CPK3-4Myc* transgenic seedlings and this phosphorylation was enhanced by osmotic stress treatment (Extended Data Fig. S2l). We failed to purify the recombinant proteins for the NB and LRR domains; therefore, we cannot rule out the possibility that CPK3 also phosphorylates other phosphosites either on SNC1^TIR^ or the NB and LRR domains.

Several other CPKs that respond to environmental stresses ^25,27,28,34^, including CPKs 4, 5, 6, 11, and 28, could also phosphorylate SNC1^TIR^ (Extended Data Fig. S2m-o), suggesting CPKs function redundantly in phosphorylation regulation of SNC1^TIR^. The CPK3-mediated phosphorylation at serines corresponding to Ser26 of SNC1^TIR^ is conserved among several plant TIRs, including the TIR domain of TIR-NLR L6 from flax (*Linum usitatissimum*) and the *Arabidopsis* TIR-only protein Response to the bacterial type III effector protein HopBA1 (RBA1) (Fig. 1j, k and Extended Data Fig. S2p). CPKs have pivotal roles in Ca^2+^ signaling in plants, ciliates and apicomplexan parasites, but not in animals or fungi; while Ca^2+^/calmodulin-dependent protein kinase II (CaMKII) resembles CPKs in animals and has a critical role in repressing SARM1-promoted axon degeneration^35^. CAMK2D could also phosphorylate SARM1^TIR^ truncated protein at the Ser567, corresponding to Ser26 of SNC1^TIR^ (Fig. 1m). These data suggest that the phosphorylation of TIRs at the conserved serine residue is conserved in plants and could be extended to animals and bacteria; however, we cannot rule out the possibility that CPK3 and CAMK2D also phosphorylate other phosphosites on TIRs and other domains.

Besides Ca^2+^ signals, BON1 also controls global osmotic stress signaling^11^. We further analyzed SNC1^TIR^ phosphorylation by other osmotic stress-activated protein kinases, such as SnRK2s and MPKs^36,37^. However, we did not find phosphorylation of SNC1^TIR^ by SnRK2.6 or MPK6 (Extended Data Fig. S3), suggesting that CPKs are major regulators of SNC1^TIR^ under osmotic stress.

### Phosphorylation of TIR represses its NADase activity

TIR NADase activity requires a conserved catalytic glutamate^1-3^. Structural analyses of SNC1^TIR^ and human SARM1^TIR^ indicate that this conserved serine (corresponding to Ser26 of SNC1^TIR^) is structurally close to the catalytic glutamate in the catalytic pocket^1,2^ (Fig. 2a,b), with a distance between the carboxylic acid group of the catalytic glutamate and the hydroxyl group of the conserved serine of 2.7 Å in SNC1^TIR^ and 2.6 Å in SARM1^TIR^ (Extended Data Fig. S4a). The phosphorylation of the conserved serines could form hydrogen bonds with NAD^+^ and repel the binding between NAD^+^ and the catalytic glutamate (Fig. 2, a, b); therefore, the phosphorylation of this conserved serine residue by CPK3 or CAMK2D may alter the catalytic pocket and thus affect TIR catalytic activity.

**Fig. 2.**
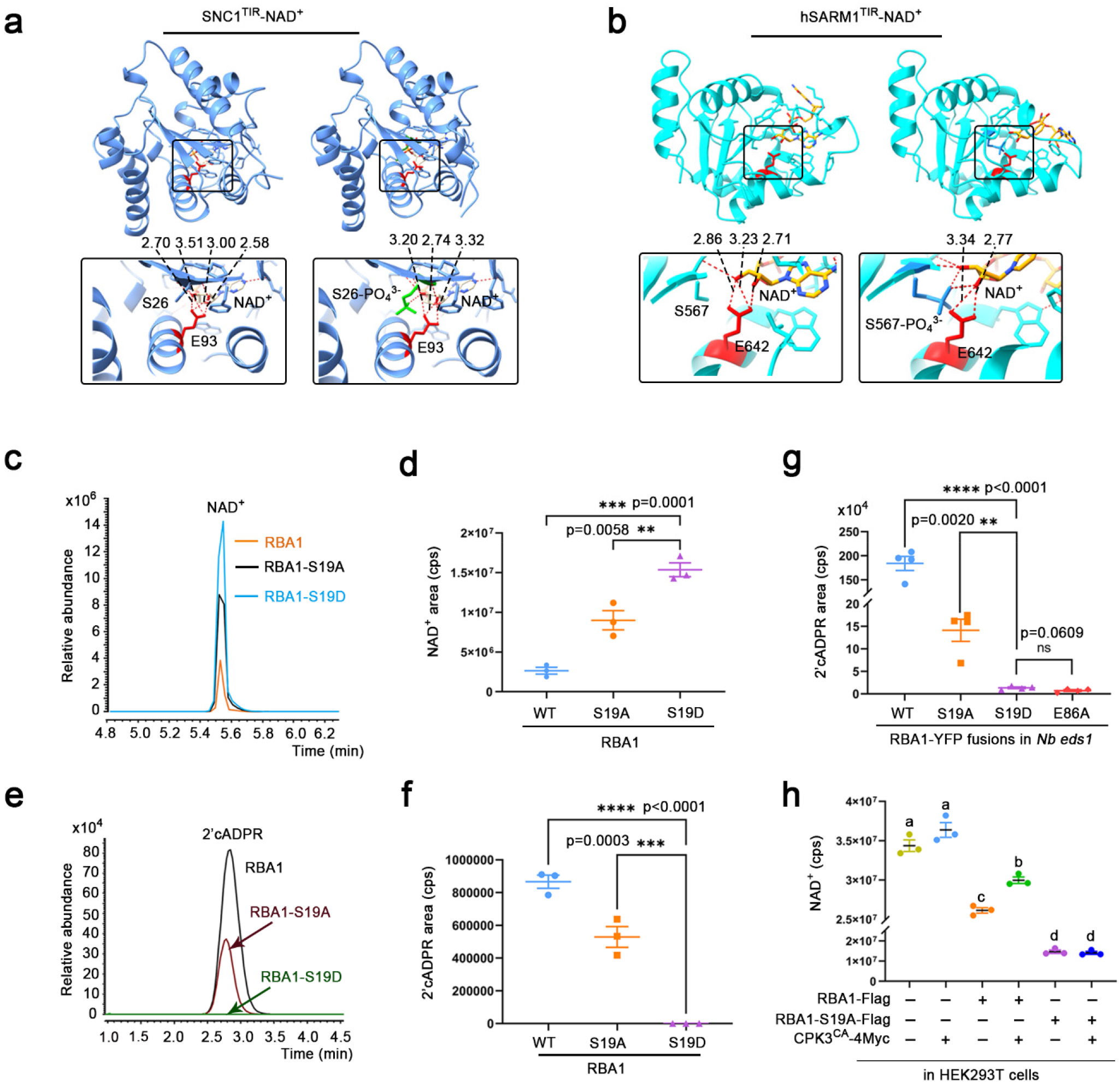
Phosphorylation of TIR represses its activity. **a**,**b**, Structures (top) and zoomed images (bottom) of SNC1^TIR^ (cornflower blue) and hSARM1^TIR^ (cyan), without (left) or with (right) modeling phosphorylation by AlphaFold3, suggest TIR phosphorylation on the conserved serine can repel the binding between the catalytic glutamate (red) in TIR domain and NAD^+^. The phosphorylated Ser26 (lime) in SNC1^TIR^ and Ser567 (dodger blue) in hSARM1^TIR^ form hydrogen bonds with NAD^+^ (wheat in **a**, and orange in **b**). Distances and numbers of hydrogen bonds between NAD^+^ and the conserved serine and glutamate are indicated in zoomed images (bottom). The sequence length of SNC1^TIR^ and hSARM1^TIR^ are the same as SNC1^TIR^ (PDB: 5TEC) and human SARM1^TIR^ (PDB: 8D0J). **c**-**f**, LC-MS/MS showing degradation of NAD^+^ (**c** and **d**) and the generation of 2′cADPR (**e** and **f**) by wild-type RBA1, RBA1-S19A, and RBA1-S19D proteins purified from insect cells *in vitro*. LC-MS/MS chromatograms (**c** and **e**) and areas under curves (**d** and **f**) indicating that RBA1-S19D could not degrade NAD^+^ or produce 2′cADPR. **g**, 2′cADPR levels in the *N. benthamiana eds1* mutant leaves transiently expressing wild-type *RBA1, RBA1-S19A, RBA1-S19D*, and *RBA1-E86A*. RBA1-S19D and -E86A could not produce 2’cADPR *in planta*. **h**, Endogenous NAD^+^ levels in the HEK293T cells expressing both wild-type and mutant *RBA1*, with or without *CPK3*^*CA*^. CPK3^CA^ represses NAD^+^ depletion mediated by wild-type but not mutant RBA1. Error bars represent means ± SEM (n = 3 experiments) for (**d, f** and **h**), and (n = 4 experiments) for (**g**). The statistical analysis was performed using a one-way ANOVA with Tukey’s test. All experiments except (**a** and **b**) were repeated at least three times with similar results.

Plant and bacterial TIRs produce two types of cyclic ADPR (cADPR) isomers^1,2,5,38,39^. One type, 3′cADPR, acts as a signaling molecule in bacterial immunity^5,38^. Plant TIRs produce several NAD^+^-breakdown products including 2′cADPR, whose function in plant immunity remains unclear but which can be monitored as a biomarker to quantify NADase activity^2^. We further analyzed the NADase activities of plant TIRs and focused on the TIR-only protein RBA1 because of its stronger *in vitro* NADase activity than the TIR domains of the TIR-NLRs^2^. Recombinant RBA1 protein purified from insect cells cleaved NAD^+^ into nicotinamide (Nam) and produced 2′cADPR *in vitro* (Fig. 2c-f, and Extended Data Fig. S4b-j). Similar to the catalytically dead E86A mutant, the phospho-mimic RBA1-S19D (corresponding to Ser26 of SNC1^TIR^) mutant abrogated the NADase activity, while the non-phosphorylatable RBA1-S19A mutant showed reduced NADase activity, as revealed by measuring the reduction of NAD^+^ and the production of 2′cADPR or Nam (Fig. 2c-f, and Extended Data Fig. S4b-j). These results show that this conserved serine residue is critical for TIR NADase activity and the non-phosphorylatable Ser-to-Ala mutation may also affect the structure and the access to the NADase catalytic site.

The repression of TIR NADase activity by phosphorylation was further investigated *in planta*. Plant TIRs require the lipase-like protein Enhanced Disease Susceptibility 1 (EDS1) for cell death and immune function^1,2,40^. The expression of plant TIR domains in the *Nb eds1* mutant resulted in the accumulation of TIR enzymatic products^2^. Consistent with the *in vitro* assays, when transiently expressed in *Nb eds1*, the catalytically dead *RBA1-E86A* and phospho-mimic *RBA1-S19D* mutant did not accumulate 2′cADPR *in planta*, while the non-phosphorylatable *RBA1-S19A* mutant produced less 2′cADPR than wild-type *RBA1* (Fig. 2g and Extended Data Fig. S4k). These results show that TIR NADase activity requires this conserved serine residue and that phosphomimetic mutation of the TIR represses its NADase activity.

The repression of plant TIR NADase activity by phosphorylation was further investigated in the immortalized human HEK293T cells. The expression of TIR-only protein RBA1 in HEK293T cells resulted in the dramatic reduction of NAD^+^ accumulation without causing cell death (Fig. 2h). To our surprise, RBA1 showed reduced NADase activity compared with that of the RBA1-S19A mutant (Fig. 2h), suggesting a putative negative regulation of RBA1 in HEK293T cells which requires this conserved serine residue. The NAD^+^ reduction by RBA1, but not RBA1-S19A, was repressed when it was transiently co-expressed with *CPK3-CA* (Fig. 2h and Extended Data Fig. S4l), suggesting that CPK3 represses plant TIR NADase activity through phosphorylation regulation in human cells.

### Phosphorylation of TIR represses its immune function and maintains plant growth

TIR NADase activity correlates with its ability to induce cell death^1,2^. All five of the studied TIRs (SNC1^TIR^, L6^TIR^, RBA1, human SARM1^tSAM-TIR^, and bacterial TIR-Ab) and the full-length SNC1 triggered cell death when transiently expressed in *Nb* leaves (Fig. 3a-h, and Extended Data Fig. S5 and S6). The induction of cell death by the plant TIRs and the full-length SNC1, but not the non-phosphorylatable SNC1^TIR-S26A^ mutant, was attenuated when they were transiently co-expressed with *CPK3-CA* (Fig. 3b-d, and Extended Data Fig. S6a-e), suggesting that the activation of CPK3 suppresses plant TIR–mediated cell death *in planta*. Like the catalytically dead TIR mutants, the phospho-mimic TIR mutants at serines corresponding to Ser26 of SNC1^TIR^, including SNC1^TIR-S26D^, SNC1^TIR-S18D/S26D (DD)^, L6^TIR-S66D^, RBA1-S19D, and hSARM1^SAM-TIR-S567D^, and the AbTIR^S140D^ mutation on the corresponding conserved serine, could not trigger cell death (Fig. 3e-h, and Extended Data Fig. S5, and S6f-l). Consistent with the NADase activity assays *in vitro* and *Nb* leaves, the non-phosphorylatable TIR mutants showed a reduced ability to trigger cell death in *Nb* leaves. In contrast, the enhanced activity of the non-phosphorylatable RBA1-S19A in HEK293T cells can be explained by unknown negative regulators of RBA1 that require this conserved serine residue (Fig. 2h). Our data demonstrate that this conserved serine residue is critical for TIR function in cell death, and the phosphorylation of TIRs at this serine suppresses their enzymatic activities and cell death phenotypes.

**Fig. 3.**
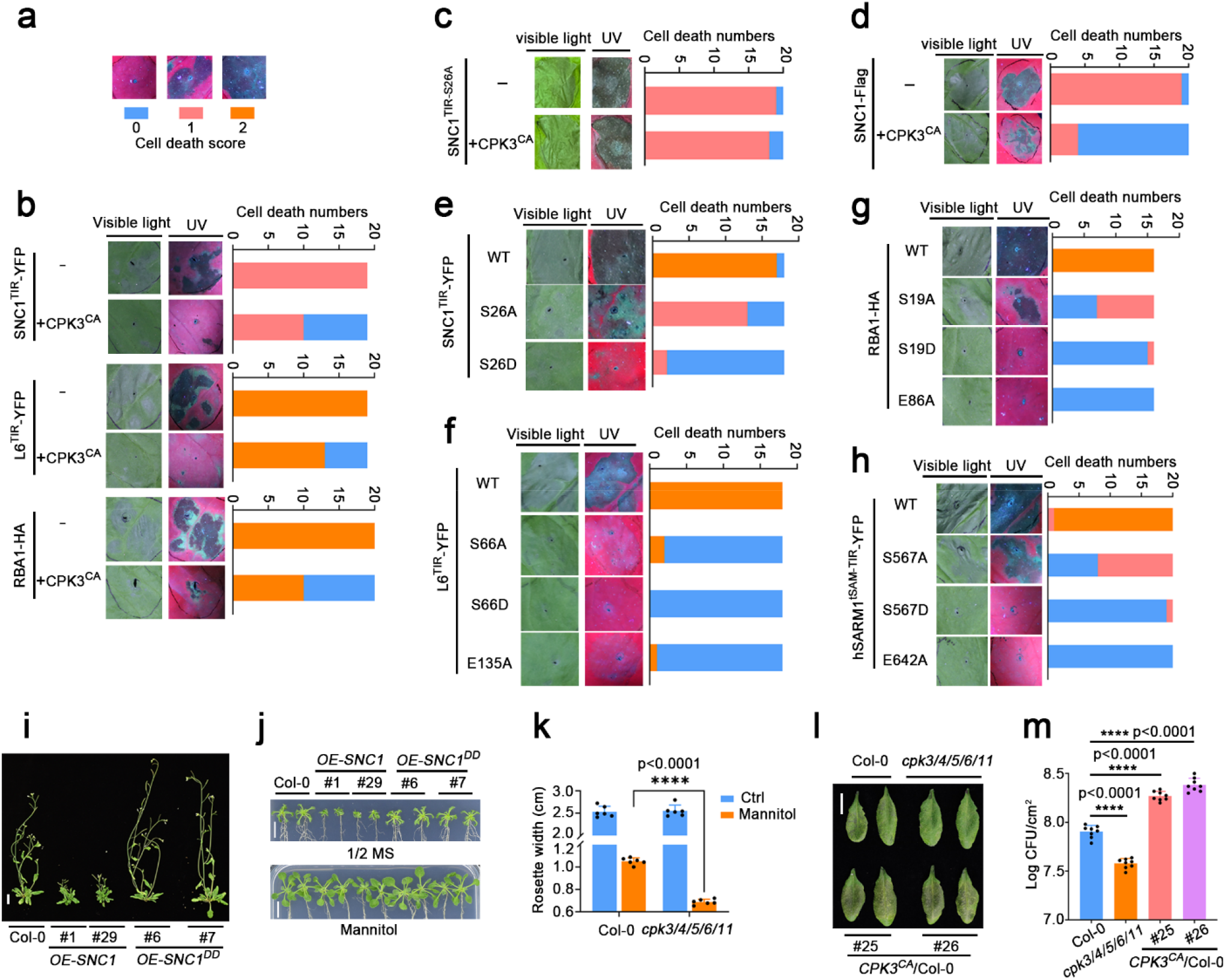
Phosphorylation of TIRs by CPKs represses TIR functions and maintains plant growth. **a**-**h**, Cell death phenotypes of wild-type and mutated TIRs *in planta*, with or without CPK3^CA^, upon transient expression in *N. benthamiana* leaves. The cell-death phenotype was visualized under visible light (left panel) or UV (right panel) and imaged 3 days after agroinfiltration inoculation. Different degrees of cell death were defined as the cell death score based on representative images (**a**). Each color on the stacked bars shows the proportions of the corresponding degrees of cell death. Constructs were driven by *35S* promoter for (**b**, and **e**-**h**), or *UBQ* promoter for (**c, d**). **b**, Expression of *SNC1*^*TIR*^, *L6*^*TIR*^, and *RBA1* with or without *CPK3*^*CA*^. **c**, Expression of *SNC1*^*TIR-S26A*^*-Flag* with or without *CPK3*^*CA*^*-HA*. **d**, Expression of full-length *SNC1-Flag* with or without *CPK3*^*CA*^*-HA*. **e**, Expression of *SNC1*^*TIR*^*-YFP*, non-phosphorylatable *SNC1*^*TIR-S26A*^*-YFP*, and phospho-mimic *SNC1*^*TIR-S26D*^*-YFP*. **f**, Expression of *L6*^*TIR*^*-YFP*, non-phosphorylatable *L6*^*TIR-S66A*^*-YFP*, phospho-mimic *L6*^*TIR-S66D*^*-YFP*, and catalytically dead *L6*^*TIR-E135A*^*-YFP*. **g**, Expression of *RBA1-HA*, non-phosphorylatable *RBA1-S19A-HA*, phospho-mimic *RBA1-S19D-HA*, and catalytically dead *RBA1-E86A-HA*. **h**, Expression of *hSARM1*^*tSAM-TIR*^*-YFP*, non-phosphorylatable *hSARM1*^*tSAM-TIR-S567A*^*-YFP*, phospho-mimic *hSARM1*^*tSAM-TIR-S567D*^*-YFP*, and catalytically dead *hSARM1*^*tSAM-TIR-E642A*^*-YFP*. **i**, Morphology of 5-week-old soil-grown plants of Col-0 and *OE-SNC1-4Myc* transgenic plants expressing wild-type or *S18D/S26D* (*DD*)-mutated *SNC1* in Col-0 background. Scale bar, 2 cm. **j**, Shoot growth of Col-0 and *OE-SNC1-4Myc* transgenic plants expressing wild-type or *S18D/S26D* (*DD*)-mutated *SNC1* in Col-0 background. The plants were transferred to and grown on the 1/2 MS medium containing 0 mM or 100 mM mannitol for 15 days. Scale bar, 0.5 cm. **k**, Rosette diameters of Col-0 and *cpk3/4/5/6/11* quintuple mutant were quantified 15 days after the seedlings were transferred to the 1/2 MS medium containing 0 mM or 100 mM mannitol. **l**,**m**, *Pst* DC3000 triggered plant immunity in Col-0, *cpk3/4/5/6/11* quintuple mutant, and *CPK3pro::CPK3*^*CA*^ in Col-0 background. Plant leaves were infiltrated with *Pst* DC3000 bacteria with OD600 at 0.002. Disease symptoms (**l**) and bacterial populations (**m**) were monitored 3 days post-infection. Scale bar, 1 cm. Error bars represent means ± SD (n = 6 seedlings) for (**k**), and (n = 8 experiments) for (**m**). The statistical analysis was performed using a one-way ANOVA with Tukey’s test. All experiments were repeated at least three times with similar results.

We further explored whether the phosphorylation of SNC1^TIR^ prevents immune signaling and maintains plant growth. Transgenic *Arabidopsis* lines overexpressing full-length *SNC1* showed autoimmune phenotypes, including dwarfism, the constitutive expression of defense genes, and elevated SA levels. By contrast, overexpression of the phospho-mimic mutant *SNC1*^*TIR-DD*^ did not trigger autoimmunity (Fig. 3i, and Extended Data Fig. S7a-c). The overexpression of wild-type *SNC1*, but not the phospho-mimic mutant, resulted in severely reduced rosette growth under osmotic stress, with elevated levels of defense marker gene expression and SA (Fig. 3j, and Extended Data Fig. S7a-c), resembling the *bon1* and *snc1-1* mutants (Extended Data Fig. S1b, l). These data demonstrate the phosphorylation of SNC1 TIR suppresses their enzymatic activities and cell death phenotypes and maintains plant growth.

We further analyzed the regulation of plant immunity by TIR phosphorylation. Consistent with the autoimmune phenotypes, lines overexpressing wild-type *SNC1* conferred resistance against *Pseudomonas syringae* (*Pst*) DC3000 compared with Col-0, while lines overexpressing the phospho-mimic *SNC1*^*TIR-DD*^ restored the susceptibility to a near wild-type (Col-0) level (Extended Data Fig. S7d-g). These data demonstrate that the phosphorylation of SNC1 TIR prevents immune signaling.

The *cpk3* mutant exhibited similar growth under hyperosmotic stress compared with Col-0 (Extended Data Fig. S8a). Since CPK3 functions redundantly with CPK4, 5, 6, 11, and 28 in TIR phosphorylation (Extended Data Fig. S2m-o), we further analyzed the phenotype of the *cpk3/4/5/6/11* quintuple mutant^41^. The *cpk3/4/5/6/11* quintuple mutant exhibited shoot growth defects and elevated levels of defense marker gene expression under hyperosmotic stress compared with Col-0 (Fig. 3k, and Extended Data Fig. S8b-d). Moreover, the *cpk3/4/5/6/11* quintuple mutant plants were resistant against *Pst* DC3000 compared with Col-0, while *CPK3pro::CPK3*^*CA*^ transgenic plants in Col-0 background were susceptible to the bacteria infection (Fig. 3l, m, and Extended Data Fig. S6m). These data demonstrate that CPKs suppress immune signaling and maintain plant growth by phosphorylation of TIRs.

### TBK1 phosphorylates SARM1 TIR and represses SARM1 TIR signaling

The phosphorylation of SARM1^TIR^ by CAMK2D at the conserved serine raises the question of whether SARM1 TIR is phosphorylated and repressed by other mammalian protein kinases. Based on genetic approaches, CaMKII, TANK binding kinase 1 (TBK1), TGFβ activated kinase (TAK1), and NIMA-related kinase 1 (NEK1) are negative regulators of axon degeneration^35,42-45^. We therefore analyzed the phosphorylation of SARM1^TIR^ by these candidate protein kinases (Fig. 4a). Besides CAMK2D, we identified the human TAK1 and TBK1 protein kinases also phosphorylate SARM1^TIR^ (Fig. 4a, b). Mutation of Ser567 to non-phosphorylatable alanines (A) reduced the phosphorylation of SARM1^TIR^ by human TBK1 and CAMK2D (Fig. 1m, 4b). The remaining phosphorylation signal of SARM1^TIR-S567A^ suggests that TBK1 and CAMK2D phosphorylation is not limited to Ser567 (Fig. 1m, 4b). Indeed, mass spectrometric analysis identified other phosphosites on SARM1^TIR^ by TBK1, and mutations of Ser574 and Ser 578 nearly abolished the phosphorylation of SARM1^TIR^ by TBK1 (Fig. 4b). These data indicate that human TBK1 phosphorylates SARM1^TIR^ on Ser567, Ser574 and Ser 578; however, whether phosphorylation of Ser574 and Ser 578 has a role in SARM1^TIR^ function requires further study.

**Fig. 4.**
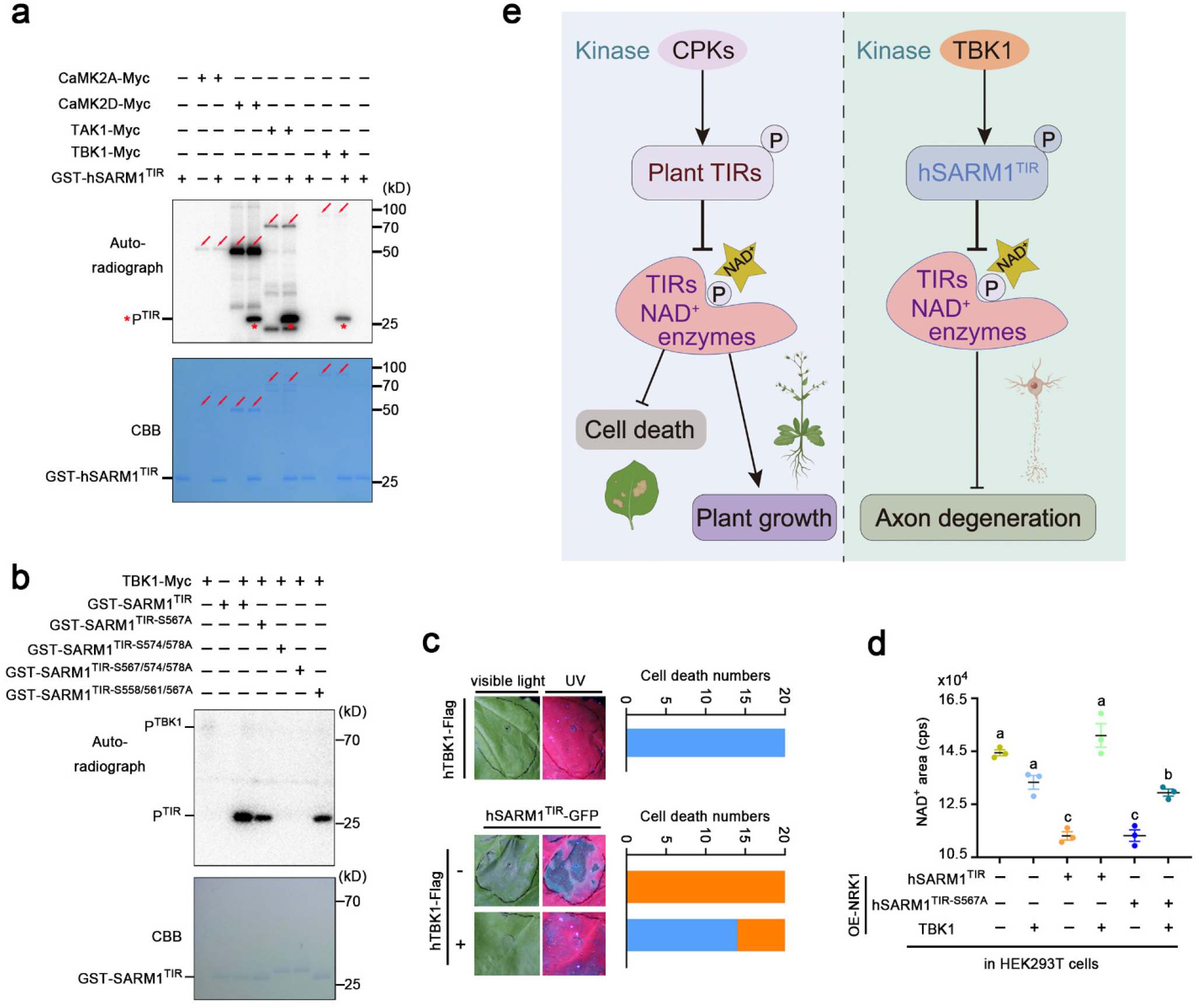
TBK1 phosphorylates SARM1^TIR^ and represses SARM1^TIR^ signaling. **a**,**b**, Phosphorylation of wild-type (**a**) and mutated (**b**) SARM1^TIR^ by CAMK2A, CAMK2D, TAK1, and TBK1. Red arrows indicated autophosphorylation of protein kinases and red asterisk indicated phosphorylation of recombinant human SARM1^TIR^ (hSARM1^TIR^). Myc-tagged human CAMK2A, CAMK2D, TAK1 and TBK1 protein kinases were immunoprecipitated from HEK293T cells expressing *CAMK2A-Myc, CAMK2D-Myc, TAK1-Myc* and *TBK1-Myc*. Autoradiography (top) and Coomassie staining (bottom) exhibit phosphorylation and the loading of CAMK2A-Myc, CAMK2D-Myc, TAK1-Myc, TBK1-Myc, and recombinant hSARM1^TIR^, hSARM1^TIR-S567A^, hSARM1^TIR-S574A/S578A^, hSARM1^TIR-S567A/S574A/S578A^, and hSARM1^TIR-S558A/S561A/S567A^. **c**, Cell death phenotypes of hSARM1^TIR^ with or without hTBK1 *in planta* upon transient expression in *N. benthamiana* leaves. The cell-death phenotype was visualized under visible light (left panel) or UV (right panel) and imaged 3 days after agroinfiltration inoculation. Different degrees of cell death were defined as the cell death score (0, blue; 2, orange). Each color on the stacked bars shows the proportions of the corresponding degrees of cell death. **d**, Endogenous NAD^+^ levels in the *NRK1*-HEK293T or *TBK1/NRK1*-HEK293T stable cell lines transiently expressing both wild-type and mutant *hSARM1*^*TIR*^. TBK1 strongly represses hSARM1^TIR^-mediated NAD^+^ depletion, but weakly represses hSARM1^TIR-S567A^-mediated NAD^+^ depletion. Error bars represent means ± SEM (n = 3 experiments). The statistical analysis was performed using a one-way ANOVA with Tukey’s test. **e**, Phosphorylation of TIR blocks TIR NADase activity and represses cell death caused by TIR overactivation to fine-regulate growth and defense trade-offs, which maintains plant growth via CPK-mediated phosphorylation of plant TIRs, and may contribute to repress axon degeneration through hSARM1 phosphorylation via CAMK2D and TBK1. TIR NADase activity is repressed by phosphorylation at a conserved serine residue spatially close to the catalytic pocket, which may repel the interaction of the catalytic glutamate with the NAD^+^ substrate. This phosphorylation-mediated inactivation of TIR represents a conserved mechanism to block TIR signaling and balance growth and defense trade-offs in plants and animals. All experiments were repeated at least three times with similar results.

SARM1^TIR^ NADase activity correlates with its function in inducing NAD^+^ depletion and cell death^1-4^. The human SARM1^tSAM-TIR^ triggered cell death when transiently expressed in *Nb* leaves. The induction of cell death by the human SARM1^tSAM-TIR^ was nearly blocked when they were transiently co-expressed with *hTBK1* (Fig. 4c and Extended Data Fig. S9a). SARM1^TIR^ triggers cell death by NAD^+^ depletion^1-4^. Nicotinamide Riboside Kinase 1 (NRK1) drives NAD^+^ biosynthesis from NAD^+^ precursors such as nicotinamide riboside (NR) in mammalian cells, which alleviates SARM1-mediated NAD^+^ depletion^3,46^. We confirmed the hTBK1-mediated repression of SARM1 TIR activity in NRK1-HEK293T cells, which enhances cellular NAD^+^ levels in the presence of NR to prevent NAD^+^ depletion-mediated cell death^3^. SARM1^tSAM-TIR^–mediated NAD^+^ depletion was completely blocked in NRK1/TBK1-HEK293T cells (Fig. 4d). Besides, human TBK1 also repressed the mutant SARM1^tSAM-TIR-S567A^–mediated NAD^+^ depletion although to a lesser extent (Fig. 4d and Extended Data Fig. S9b), indicating TBK1 represses SARM1 TIR through phosphorylation of Ser567 and other sites. Moreover, previous genetic studies in *Caenorhabditis elegans* indicated a pivotal role of CaMKII in suppressing SARM1-MAPK-promoted axon degeneration^35^. These data suggest that TBK1 and CaMKII suppress SARM1 TIR signaling through direct phosphorylation.

## DISCUSSION

Together, we discovered that CPK, CAMK2D and TBK1 protein kinases block TIR immune signaling through phosphorylation of a conserved serine spatially close to the catalytic glutamate in TIR domains in plants and animals, respectively (Fig. 4e). This phosphorylation abolishes the TIR NADase enzymatic activity, represses plant immunity, and subsequently confers growth maintenance under osmotic stress or in soil indirectly. In addition, the TBK1 protein kinase-mediated TIR phosphorylation represses SARM1-mediated NAD^+^ depletion. Thus, TIR phosphorylation represents a conserved mechanism to block TIR signaling and balance growth and defense trade-offs in plants and animals.

Ca^2+^ signals are transiently triggered by multiple stimuli. The plasma membrane-localized BON proteins directly interact with SOMATIC EMBRYOGENESIS RECEPTOR-LIKE KINASEs (SERKs) and Ca^2+^ pumps in plants^23,47,48^, and may mediate Ca^2+^ signals and other downstream responses during plant development, brassinosteroid (BR), immune, and osmotic stress signaling^11,20,23^, probably via affecting the microdomain formation during cell surfacing signaling^47^. Most of the CPKs such as CPK3 are activated by Ca^2+^ signals and mediate stress responses to environmental changes and this process may require proper function of BON proteins in cell surface signaling; some of the CPKs are almost Ca^2+^-independent and probably contribute to basal CPK signaling^26^. CPK-mediated phosphorylation of TIR is not limited to CPK3 and is likely to suppress TIR-NLR signaling to maintain plant growth and ensure abiotic stress responses and may also be involved in buffering plant immunity^11,49,50^; however, the spatial and temporal specificity of CPKs in plant responses to stimuli requires further investigation. Therefore, adequate TIR phosphorylation in plants may require the coordination of BON-mediated cell surface signaling, Ca^2+^ signals, and the Ca^2+^-dependent CPK functions, and our findings explain how BON represses TIR immune signaling via mediating Ca^2+^ signaling. It will be interesting to determine how CPKs are regulated during effector-triggered immunity and whether BON proteins are indeed required for CPK activation triggered by cell surface signaling. Plant TIR signaling is activated in a two-tiered manner: pathogen-dependent initial activation in infected cells, and the subsequent transcription of many *TIR*-encoding genes in infected cells and neighboring uninfected cells^39,51,52^. TIR phosphorylation may play a critical role in the attenuation of immunity after the successful control of pathogen infection, especially in neighboring uninfected cells. Our findings thus provide a plausible explanation for the balance and switch between stress responses and plant growth.

SARM1 is a central executioner of axonal degeneration and triggers SARM1-dependent cell death^53^. Genetic analyses of SARM1 mutants suggest that loss of SARM1 can be an attractive therapeutic strategy for neurodegenerative diseases^39^. TBK1, TAK1 and CaMK2D directly phosphorylate SARM1 TIR and TBK1 represses NAD^+^ depletion when expressed in NRK1-HEK293T cells (Fig. 4). CaMKII has a pivotal role in repressing SARM1-promoted axon degeneration^35^; however, whether CAMK2D- and TBK1-mediated phosphorylation and repression of SARM1 occur during axon degeneration requires further study. Our discovery of CAMK2D- and TBK1-mediated phosphorylation of SARM1 and repression of SARM1 activity provide an explanation for their function in repressing axon degeneration.

## EXPERIMENTAL PROCEDURES

Further details and an outline of resources used in this work are provided in the Extended Data Methods, including plant materials and growth conditions, cloning the *ios* mutants, plasmid construction, agrobacterium-mediated transient expression, SARM1 transiently expressed in *NRK1*- or *TBK1/NRK1*-HEK293T stable cell line, recombinant protein expression and purification, *in vitro* kinase assays, immunoprecipitated kinase assay, generation of anti-phosphorylation antibodies and immunoblotting, data-independent acquisition (DIA) based mass spectrometry assay, *in vitro* NADase assay, endogenous mammalian cell NAD^+^ quantification, LC-MS/MS metabolite measurement, bacterial disease assays, SYPRO ruby gel staining, co-immunoprecipitation assay, split luciferase (LUC) complementation assay, RNA extraction and qRT-PCR, and Ca^2+^ signal detection with aequorin-based Ca^2+^ reporter in plants.

## QUANTIFICATION AND STATISTICAL ANALYSIS

The statistical analysis was performed using a one-way ANOVA with Tukey’s test, a two-way ANOVA with Tukey’s test, or Student’s t-test, in assays related to gene expression, plant growth, and SA and metabolites quantifications.

## ACKNOWLEDGMENTS

We thank Jeffery. L. Dangl and Marc. T. Nishimura for careful reading and discussion of the manuscript. We thank the members of the Zhao Lab for helpful discussions. This work was supported by the Strategic Priority Research Program of the Chinese Academy of Sciences grants XDB063020102 (to Y.Z.), and XDB27040214 (to L.W.), and the Science and Technology Commission of Shanghai Municipality grant 22ZR1481400 (to Y.Z.). Y.Z. was supported by the Shanghai Center for Plant Stress Biology, Chinese Academy of Sciences, and Key Laboratory of Plant Carbon Capture, Chinese Academy of Sciences. L.W. was supported by the National Key Laboratory of Plant Molecular Genetics, the Institute of Plant Physiology and Ecology/Center for Excellence in Molecular Plant Sciences.

## SUPPLEMENTAL INFORMATION

Supplemental Information includes Extended Data Methods and nine figures.

## AUTHOR CONTRIBUTIONS

Y.Z. and L.W. conceived and designed the research. J.L., S.C., B.Y., Q.L., R.L. and Z.W. performed the experiments. J.L., S.C., B.Y., L.W., and Y.Z. analyzed the results. Y.Z., J.L. and L.W. wrote the manuscript.

