## Supplemental information for "TIR immune signaling is blocked by phosphorylation to maintain growth in plants and animals"

### Extended Data Methods

#### Plant materials

Arabidopsis wild-type Col-0, *AEQsig6*[1], *bon1-1*[1], *bon1-7*[1], *snc1-1*[2], *cpk3-2* and *cpk3/4/5/6/11* quintuple mutants were used[3]. The *ios* mutants were isolated from an ethyl methanesulfonate (EMS)-mutagenized mutant pool using the *bon1-7* mutant as wild-type. To generate wild-type and mutated (*G1343D*, *V518I*, and *S18DS26D*) *OE-SNC1-4Myc* transgenic plants, the genomic sequences (without stop codon) were cloned into pCAMBIA1300, confirmed by sequencing, and transformed into the Col-0, *bon1-1* or *bon1-7* mutant using the Agrobacterium-mediated floral dip method. To generate wild-type and $CPK3^{CA}$, $CPK3^{CA\text{-}K107M}$ and $CPK3^{CA\text{-}\Delta K107}$ mutated *CPK3proCPK3-4Myc* transgenic plants, the *CPK3* genomic sequences (without stop codon), $CPK3^{CA}$, $CPK3^{CA\text{-}K107M}$ and $CPK3^{CA\text{-}\Delta K107}$ sequences were cloned into pCAMBIA1300[4], confirmed by sequencing and transformed into the Col-0 and *bon1-1*. *Nicotiana benthamiana* wild-type and *eds1-2* mutant were used[5].

#### Plant growth conditions

In most cases, the Arabidopsis plants were grown under a long photoperiod (16 h light, 23°C; 8 h dark, 23°C) with 60%-70% relative humidity. The seeds were sterilized with 5% sodium hypochlorite for 10 minutes and rinsed with sterile-deionized water for 5 times. Seeds were grown on 1.2% agar containing 1/2 MS nutrients (PhytoTech, M542), 1% sucrose, and pH 5.7. Plates were stratified at 4°C for 3 days. For the shoot growth phenotype assay, 3-day-old vertically grown seedlings were transferred from 1/2 MS medium to medium containing 0 mM or 100 mM mannitol for 15 days. For SA measurement and transcriptional analysis, 3-day-old vertically grown seedlings were transferred from 1/2 MS medium to medium containing 0 mM or 100 mM mannitol for 5 days. The seedlings were grown on medium in a Percival CU36L5 incubator at 23°C under long-day conditions (16 h light/8 h dark).

For *Nicotiana benthamiana*, the plants were grown in a phytotron maintained at a photoperiod (12 h light, 26°C; 12 h dark, 22°C). The 4-5 weeks *Nicotiana benthamiana* leaves were used for *Agrobacterium tumefaciens*-mediated transient expression of *TIRs* and *CPK3* for the hypersensitivity reactions, split-luciferase, and co-immunoprecipitation assays.

#### Cloning the *ios* mutants

The genomes of *bon1-7* (wild-type) and the *ios* mutants were sequenced, and used as reference genomes to identify genetic markers. To perform the genetic analysis, the *ios* mutants were backcrossed with *bon1-7* to construct the $F_2$ populations. Since these *ios* mutants are dominant mutations, genomic DNA was extracted from 50 individual $F_2$ seedlings showing the *bon1*-like phenotypes, and mixed in an equal ratio. At least 6 μg of bulked DNA was used to construct a sequencing library according to the manufacturer's instructions (Illumina Inc.). The 400 bp DNA paired-end library was sequenced using the Illumina HiSeq system (PE150) at Shanghai Majorbio Bio-pharm Technology Co., Ltd. Sliding window methods were used to determine the SNP/InDel index of the whole genome. The difference between the SNP/InDel index of pools was calculated as ΔSNP/InDel index. Finally, the mutation area was defined based on the linkage of genetic markers between the $F_2$ seedling library and the *bon1-7* reference genome.

**Plasmid construction for plant transformation**

The coding sequences of *RBA1* (residues 1-191) and *RBA1* mutants (*S19A*, *S19D*, and *E86A*)[6], $SNC1^{TIR}$ (residues 10-235) and $SNC1^{TIR}$ mutants (*S18A*, *S18D*, *S26A*, *S26D*, *S18AS26A*, *S18DS26D*, and *E93A*), $L6^{TIR}$ (residues 1-233) and $L6^{TIR}$ mutants (*S66A*, *S66D*, and *E135A*), $hSARM1^{tSAM\text{-}TIR}$ (residues 409-724) and mutants (*S567A*, *S567D*, and *E642A*)[7], were amplified by PCR and cloned into pENTR and pUC19 vectors that contained the recombination sites of *attL1* and *attL2* arms for LR reaction, and then recombined into the Gateway destination vector pEarley (fused to YFP or HA).

The coding sequences of $CPK3^{CA}$, $CPK3^{CA\text{-}K107M}$ or the genomic sequences of *SNC1* were amplified by PCR and cloned into *pCAMBIA1305-UBQ:HA* or *pCAMBIA1305-UBQ:Flag* vectors and used for plant transformation.

**Agrobacterium-mediated transient expression**

For agroinfiltration in *Nicotiana benthamiana*, the *Agrobacterium tumefaciens* strain GV3101 was grown overnight. Cells were harvested by centrifugation and re-suspended in the infiltration medium (10 mM $MgCl_2$, 10 mM MES, pH 5.6, and 150 μM acetosyringone), with OD600 nm at indicated levels. The Agrobacterium carrying *35S:P19*, a viral suppressor of gene silencing, was co-injected with OD600 nm at 0.1 to avoid transgene-triggered gene silencing. The cell-death images were taken at 3-5 days post-infection (dpi) under visible light or UV conditions. For western blot analysis, the infected leaf samples were collected at 2 dpi. The transformed leaves were collected at 2-3 dpi for NADase metabolite assay. Four leaf disks (9 mm in diameter) collected at 4 leaves from 3 different plants were snap-frozen and homogenized into powder. The samples were dissolved in 200 μL 50% (v/v) methanol and vortexed for 2 minutes. Plant extracts in 50% methanol were subsequently centrifuged and stored at -80°C until LC-MS/MS analysis.

**SARM1 transiently expressed in *NRK1*- or *TBK1/NRK1*-HEK293T stable cell line**

A HEK293T cell line stably expressing NRK1 was constructed as previously described[7]. For generating lentivirus, the HEK293T cells were seeded 24 hours before transfection. Plasmids containing TBK1-4×Myc or vector along with packaging plasmids PSPAX and PMD2.G were transfected using Lipofectamine 2000 (Invitrogen) to generate lentivirus. The supernatants containing infectious lentiviruses were collected after 24 h and 48 h, and filtered through 0.45 µm filters (Millipore), then frozen at -80°C. To generate the *TBK1/NRK1*-HEK293T stable cell line, *NRK1*-HEK293T stable cells were transduced with the TBK1-4×MYC lentivirus. The efficiency of TBK1 overexpression was determined by Western blot. Cells were infected with the control vector as described above. The *NRK1/TBK1*-HEK293T or *NRK1/Control*-HEK293T cells were seeded into a 6-well plate 24 h before transfection. Cells were transfected with Lipofectamine 2000 (Invitrogen) mixed with a plasmid containing 3×FLAG-SARM1$^{tSAM\text{-}TIR}$ or the plasmid containing 3×FLAG (containing the starting codon) as the control. Nicotinamide Riboside (NR) was added at a final concentration of 1 mM to improve cell viability. After 2 days the cells were harvested for further research.

**Recombinant protein expression and purification**

The *RBA1* (residues 1-191) and *RBA1* mutants (*S19A*, *S19D*) coding sequences were cloned into the pFastBac 1 plasmid (Invitrogen) with an N-terminal Flag tag and C-terminal 6×His tag[6]. The constructs generated recombinant baculovirus in Sf9 insect cells (Invitrogen). RBA1 and RBA1 mutant proteins were expressed in Sf9 insect cells with recombinant baculovirus infection at 28°C and 100 rev min$^{-1}$ for 96 h. The cells were collected by centrifugation and lysed by sonification in lysis buffer composed of 50 mM Tris–HCl pH 7.5, 300 mM NaCl, 10 % glycerol, 5 mM ATP, 1 mM EDTA, 0.2 % chaps and 1×Protease Inhibitor Cocktail. Cell debris was centrifuged at 20 000 g for 1 h at 4°C. The supernatant was filtered and incubated with anti-Flag G1 affinity resin (GenScript) at 4°C for 3 h. After washing with ten-column volumes of lysis buffer, the resins containing RBA1 proteins were used for *in vitro* NADase assays.

The *SNC1*$^{TIR}$ (residues 1-42), *SNC1*$^{TIR}$ mutants (*S18A*, *S26A* and *S18AS26A*), *CPK3*, *CPK4*, *CPK5*, *CPK6*, *CPK11* and *CPK28* were cloned into pGEX4T-1 vector with termination codon and expressed in

E. *coli* with N-terminal GST tag. The construction vectors were transformed into E. *coli* BL21 (DE3) and the cell cultures were grown at 37°C to OD600 nm at 0.6-0.8. Isopropyl-β-D-thiogalactoside (IPTG) was added to induce protein expression at 18°C for 16 h before harvesting. The cells were lysed using sonication in lysis buffer composed of 20 mM Tris–HCl, pH 8.0, 200 mM NaCl and 5 mM DTT. The resulting supernatant was incubated with Glutathione Sepharose 4B resin at 4°C for 3 h. After binding to the Glutathione agarose, the agarose was washed with ten-column volumes of lysis buffer. Bound protein was eluted using 20 mM Tris–HCl, pH 8.0, 200 mM NaCl and 10 mM reduced glutathione. $L6^{TIR}$ (residues 27-229) and $L6^{TIR}$ mutant (*S66A*), were inserted into the $pET30a^{+}$ vector with termination codon using an N-terminal 6×His tag. Plasmids were expressed in E. *coli* BL21 (DE3) and induced according to the auto-induction method[8]. $L6^{TIR}$ and mutants were as previously described[9]. $hSARM1^{TSAM-TIR}$, as previously described[10], was codon-optimized for *E. coli* expression using the Genscript codon optimization tool, amplified and cloned into $pET30a^{+}$ or pGEX4T-1. The construct was transformed into Shuffle T7 Express Competent *E. coli* (NEB) ($pET30a^{+}$-$hSARM1^{TSAM-TIR}$) or E. *coli* BL21 (DE3) (*pGEX4T-1-*$hSARM1^{TSAM-TIR}$).

### ***In vitro*** **kinase assays**

Recombinant GST-CPK3, GST-CPK4, GST-CPK5, GST-CPK6, GST-CPK11, GST-CPK28, TBK1-Myc (Leading Biology, PH39446M5), CAMK2A-Myc (Leading Biology, PH35249M5), CAMK2D-Myc (Leading Biology, PH35097M5) and TAK1-Myc (Leading Biology, PH39801M5) were incubated with recombinant GST-$SNC1^{TIR}$, GST-$SNC1^{TIR-S18A}$, GST-$SNC1^{TIR-S26A}$, GST-$SNC1^{TIR-S18AS26A\ (AA)}$, $L6^{TIR}$-His, $L6^{TIR-S66A}$-His, GST-$hSARM1^{TIR}$, GST-$hSARM1^{TIR-S567A}$, GST-$hSARM1^{TIR-S574A/S578A}$, GST-$hSARM1^{TIR-S567A/S574A/S578A}$, GST-$hSARM1^{TIR-S558A/S561A/S567A}$, $hSARM1^{TIR}$-His, and $hSARM1^{TIR-S567A}$-His in reaction buffer (25 mM Tris–HCl, pH 7.5, 10 mM $MgCl_2$, 0.25 mM DTT, 1 mM ATP, 2 μl Ci [γ-$^{32}P$] ATP) for a total reaction volume of 20 μL at 30°C. After incubation for 3h, the proteins were separated by SDS–PAGE and stained with Coomassie blue. After electrophoresis, the gel was dried under a vacuum at 80°C for 1.5 h on filter paper and then exposed to a phosphor imager overnight. Phosphorylation was visualized by a Personal Molecular Imager.

For *in vitro* phosphorylation assays followed by PNBM alkylation[11], recombinant GST-CPK3, MBP-SnRK2.6, MBP-MAPK6, and MBP-$MKK4^{DD}$, were incubated with GST-$SNC1^{TIR}$, GST-$SNC1^{TIR-S18A}$, GST-$SNC1^{TIR-S26A}$, GST-$SNC1^{TIR-S18AS26A\ (AA)}$, RBA1-Flag, and RBA1-S19A-Flag, in reaction buffer containing 25 mM Tris–HCl, pH 7.5, 10 mM $MgCl_2$, 0.25 mM DTT, 0.5 mM ATPγS (Abcam, ab138911) at 30°C for 1 h. Then, PNBM (Abcam, ab138910) in DMSO was added to afford a final concentration of 2.5 mM with 5% DMSO. After incubation for 1 h, the proteins were separated by SDS–PAGE and then transferred to PVDF membrane for immunoblot detection. The anti-thiophosphate ester antibody (Abcam, ab92570) was used to detect phosphorylated proteins. Anti-GST antibody (Thermofisher, Cat#: 136700) was used to detect the GST-CPK3 and wild-type and mutated GST-SNC1$^{TIR}$. The anti-Flag antibody (ABclonal, AE024) was used to detect the RBA1-Flag and RBA1S19A-Flag.

**Immunoprecipitated kinase assay**

The seven-day-old *CPK3proCPK3-4Myc* transgenic seedlings in Col-0 background, grown on 1/2 MS plate, were exposed to 1/2 MS medium containing 0 or 0.6 M mannitol for 30 mins. Immunoprecipitated kinase assays were also performed using the *Nicotiana benthamiana* leaves transiently expressed *UBQ:CPK3$^{CA}$-HA* or *UBQ:CPK3$^{CA-K107M}$-HA*. Total proteins were extracted with immunoprecipitation buffer containing 20 mM Tris–HCl (pH 7.6), 150 mM NaCl, 5 mM $MgCl_2$, 10 mM $Na_3VO_4$, 10 mM NaF, 50 mM β-Glycerophosphate disodium, 0.5 mM DTT, 10 mg ml$^{-1}$ leupeptin, 10 mg ml$^{-1}$ antipain, 10 mg ml$^{-1}$ aprotinin, 0.1% NP-40 and 10% glycerol, and then incubated with anti-Myc or anti-HA magnetic beads for 3 h at 4°C. After incubation, the beads were washed three times with wash buffer (20 mM Tris–HCl, pH 7.6, 150 mM NaCl) and three times with kinase buffer (25 mM Tris–HCl, pH 7.5, 10 mM $MgCl_2$, 0.25 mM DTT), and then performed the phosphorylation assay as stated above.

**Generation of anti-phosphorylation antibodies and immunoblotting**

Anti-phosphorylation rabbit antibodies were generated by ABclonal Technology. The HPLC high-purity modified peptide C-DVFP(S-p)FRGED was used as an antigen to generate polyclonal anti-phosphorylation antibodies for S26 in SNC1. The phosphorylation site-specific antibodies were purified using the non-modified peptides C-DVFPSFRGED, which absorbed the non-specific antibodies. The anti-phosphorylation antibodies for S26 in SNC1 can detect Ser26 phosphorylation with recombinant SNC1$^{TIR}$ protein and immunoprecipitated SNC1$^{TIR}$ and full-length SNC1 in plants.

Flag-tagged SNC1$^{TIR}$ or full-length SNC1 was immunoprecipitated from *Nicotiana benthamiana* leaves at 48 hours after transiently expressing *UBQ:SNC1$^{TIR}$-Flag* or full-length *UBQ:SNC1-Flag*, with or without *UBQ:CPK3$^{CA}$-HA* or *UBQ:CPK3-HA*, with or without 1 M mannitol treatment for 2 hours. Total proteins were extracted with immunoprecipitation buffer containing 20 mM Tris–HCl (pH 7.6),

150 mM NaCl, 5 mM $MgCl_2$, 10 mM $Na_3VO_4$, 10 mM NaF, 50 mM β-Glycerophosphate disodium, 0.5 mM DTT, 10 mg $ml^{-1}$ leupeptin, 10 mg $ml^{-1}$ antipain, 10 mg $ml^{-1}$ aprotinin, 0.1% NP-40 and 10% glycerol, and then incubated with anti-Flag magnetic beads for 3 h at 4°C. After incubation, the beads were washed three times with wash buffer containing 20 mM Tris–HCl (pH 7.6) and 150 mM NaCl, and then treated with or without λ-protein phosphatase (λ-PPase) for 30 minutes at 30°C. For western blots, the membranes were incubated at 4°C overnight in TBST with 5% skim milk containing 1:100 diluted anti-phospho-Ser26-SNC1$^{TIR}$ antibody, and then washed three times (10 min each) with TBST. The membranes were then incubated for 3 h at room temperature in TBST with 3% skim milk containing 1:5000 diluted goat anti-rabbit antibody, and then washed five times (5 min each) with TBST and detected using e-BLOT Touch Imager (e-BLOT Life Science, Shanghai) or Tanon-5200 Chemiluminescent imaging system (Tanon).

**Data-independent acquisition (DIA) based mass spectrometry assay**

Flag-tagged SNC1$^{TIR}$ was immunoprecipitated from *Nicotiana benthamiana* leaves at 48 hours after transiently expressing *UBQ:SNC1$^{TIR}$-Flag* and *UBQ:CPK3$^{CA}$-HA*. The SNC1$^{TIR}$-Flag protein was extracted with anti-Flag magnetic beads using SDT lysis buffer (4% SDS, 100 mM DTT, 100 mM Tris–HCl, pH 8.0). The SNC1$^{TIR}$-Flag magnetic beads were boiled for 3 min and further ultrasonicated. The undissolved beads were removed by centrifugation at 16 000 g for 15 min. Proteins in the supernatant were collected. Protein digestion was performed using filter-aided sample preparation (FASP) method, with trypsin overnight at 37°C. DTT and iodoacetamide were added to the UA buffer. The peptides were collected by centrifugation at 16 000 g for 15 minutes and desalted with C18 StageTip for further LC-MS analysis. LC-MS/MS was performed on an Orbitrap Astral mass spectrometer coupled with a Vanquish Neo UHPLC system (Thermo Fisher Scientific). The DIA method consisted of a survey scan from 380-980 m/z at resolution 240000 with an AGC target of 500% and 5ms injection time. The DIA MS/MS scans were acquired by Astral from 150-2000 m/z with a 2 m/z isolation window and with an AGC target of 500% and 3ms injection time. The normalized collision energy was set to 25, and the cycle time was set to 0.6s. The spectra of the full MS scan and DIA scan were recorded in profile and centroid type, respectively, and then the DIA MS data were analyzed using Spectronaut 18 (Biognosys AG, Switzerland).

***In vitro* NADase assay**

Ten microliters of anti-flag beads laden with purified protein were incubated with 30 μM $NAD^+$ (final concentration) and reaction buffer (92.4 mM NaCl and 0.64 × PBS), for a total reaction volume of 50 μL. Reactions were carried out at room temperature (25°C) for 2 h, and stopped by the addition of 50 μL of 1 M of perchloric acid ($HClO_4$) and placing the tube on ice for at least 10 min. Neutralization was performed with 16.7 μL 3 M $K_2CO_3$. Samples were placed on ice for 10 min, and then separated by centrifugation. For LC-MS/MS analysis, the extraction was performed using 50% methanol in distilled water (see LC-MS/MS metabolite measurement for further details).

**Endogenous mammalian cell $NAD^+$ quantification**

HEK293T and NRK1-HEK293T cultures were harvested at 3 000 rpm for 1 min at 25°C, then lysed by adding methanol: acetonitrile: water (2: 2: 1). The cells were mixed and freeze-thawed by liquid nitrogen for three times. To precipitate the proteins, samples were deposited at -20°C for one hour and then centrifuged at 12000 rpm at 4°C for 15 min. The supernatants were dried with a Termovap sample concentrator and redissolved with 100 μL of 50% methanol in distilled water. The sample was stored at -80°C for LC-MS/MS analysis.

**LC-MS/MS metabolite measurement**

Samples were prepared by mixing the reactions with 50% methanol in distilled water. The samples were centrifuged and applied to the LC-MS/MS for metabolite identification and quantification. Samples were analyzed by Q Exactive quadrupole orbitrap high-resolution mass spectrometry coupled with a Dionex Ultimate 3000 RSLC (HPG) ultra-performance liquid chromatography (UPLC-Q-Orbitrap-HRMS) system (Thermo Fisher Scientific), with a HESI ionization source under positive ion modes using a heated ESI source. Samples were separated with an ACQUITY UPLC HSS T3 column (100 mm × 2.1 mm, 1.8-μm particle size; Waters). The mobile phase consisted of 2 mM ammonium formate in water (A), and 100% acetonitrile (B); the gradient elution was set as follows: 0–2.00 min, 1% B; 2.00–7.00 min, 1%–95% B; 9.00–9.10 min, 95%–1% B; 9.10–11.00 min, 1% B. The flow rate was set to 0.3 mL/min and column temperature at 45°C. Metabolites were quantified by using areas, and the retention time for each compound was determined with standard compounds, including $NAD^+$, cADPR, and Nam, reconstituted with 50% methanol in distilled water. The metabolites were also detected and quantified with QTRAP 6500+ mass spectrometer (AB SCIEX) under positive ESI multiple reaction monitoring (MRM) for monitoring analyte parent ion and product ion formation. MRM conditions were optimized using authentic standard chemicals including: NAD+ ([M+H]+ 664 > 136.00, 664 > 428, 664 > 542); Nam ([M+H]+ 123 > 80); cADPR ([M+H]+ 542 > 136, 542 > 348, 542 >428); v-cADPR ([M+H]+ 542 >136).

**Bacterial disease assays**

For *Pst* DC3000 bacterial inoculation, *Pst* DC3000 strains were incubated overnight in Luria–Bertani (LM) medium at 30°C to an OD600 of 0.8–1.0, as previously described[12]. Bacteria were collected by centrifugation and washed once with sterile water, and adjusted to a cell density of OD600 nm of 0.2 ($1x10^8$ cfu/mL). Before bacteria infiltrated into leaves, bacterial suspension was further diluted to a cell density of about OD600 nm of 0.002. Bacteria were infiltrated into leaves with a needleless syringe. Inoculated plants were kept under ambient humidity for 2 h to allow evaporation of excess water from the leaf and then covered with a transparent plastic dome to maintain high humidity. For the quantification of bacteria, four leaf discs from two different leaves (after surface sterilization with 75% ethanol) were taken using a cork borer (7 mm in diameter) as one biological repeat, and three repeats were taken for each treatment. Leaf discs were ground and diluted in sterile water, and the extraction solutions were then plated on LM agar plates supplemented with rifampicin (at 50 mg/L). Colonies were counted with a stereoscope 24 h after incubation at 30°C. For the SA measurement assay, after bacteria were infiltrated into leaves for three days, the leaves were collected, and the SA levels were detected by HPLC.

**SYPRO ruby gel staining**

Total proteins were boiled in loading buffer composed of 50 mM Tris–HCl, pH 6.8, 2% SDS, 10% (v/v) glycerol, 0.1% (w/v) bromophenol blue, and 1 mM DTT for 10 min and separated by SDS–PAGE. After electrophoresis, the gel was transferred to a nitrocellulose membrane and incubated in the SYPRO Ruby Protein Gel stain. The membrane was washed with 1% acetic acid solution for 30 min, rinsed in distilled water for 10 minutes, and taken photos.

**Co-immunoprecipitation assay**

The *35S:SNC1*$^{TIR}$*-Myc* construct was co-expressed with *35S:CPK3-YFP* or *35S:YFP-YFP* in *eds1* mutant leaves of *Nicotiana benthamiana*. At two days post-infection, total proteins were extracted with immunoprecipitation buffer containing 20 mM Tris–HCl (pH 7.5), 150 mM NaCl, 5 mM DTT, 0.5%

NP40 and 1×protease inhibitor cocktail (Thermo Fisher, Cat#: 78442), followed by incubating with anti-GFP beads for 3 h, then the beads were washed with immunoprecipitation buffer for 5 times and with 1×PBS for 3 times. Immunoblot analysis was performed using anti-GFP or anti-Myc antibodies (Shanghai Epizyme, Cat#: LF302).

The *UBQ:SNC1$^{TIR}$-Flag* construct was co-expressed with *UBQ:CPK3$^{CA}$-HA* or *UBQ:CPK3$^{CA-K107M}$-HA* in *eds1* mutant leaves of *Nicotiana benthamiana*. At two days post-infection, total proteins were extracted with immunoprecipitation buffer containing 20 mM Tris–HCl (pH 7.6), 150 mM NaCl, 5 mM $MgCl_2$, 10 mM $Na_3VO_4$, 10 mM NaF, 50 mM β-Glycerophosphate disodium, 0.5 mM DTT, 10 mg $ml^{-1}$ leupeptin, 10 mg $ml^{-1}$ antipain, 10 mg $ml^{-1}$ aprotinin, 0.1% NP-40 and 10% glycerol, and then incubated with anti-HA magnetic beads for 3 h at 4°C. After incubation, the beads were washed three times with wash buffer containing 20 mM Tris–HCl (pH 7.6) and 150 mM NaCl. Immunoblot analysis was performed using the anti-HA antibody (Roche, Cat#: 11867423001) or anti-Flag antibody (ABclonal, AE024).

**Split luciferase (LUC) complementation assay**

The coding sequence of *SNC1$^{TIR}$* and *CPK3* were cloned into pCAMBIA1-cLUC and pCAMBIA1-nLUC, respectively. The constructs were co-transformed into the *eds1* mutant leaves of *Nicotiana benthamiana*. At two days post-infection, LUC signals were detected with a CCD camera (Tanon).

**RNA extraction and qRT-PCR**

Total RNA was extracted from the seedlings five days after being transferred to the 1/2 MS medium containing 0 mM or 100 mM mannitol. Approximately 1 μg of total RNA was used for first-strand cDNA synthesis, with the HiScript ® 1st Strand cDNA Synthesis Kit (Vazyme). The cDNA products were used for real-time PCR analysis using the 2×Universal SYBR Green Fast qPCR (ABclonal, Wuhan, China). The data were recorded and analyzed using the CFX96 Touch real-time PCR detection system (Bio-Rad).

**$Ca^{2+}$ signal detection with aequorin-based $Ca^{2+}$ reporter in plants**

The *bon1-7* and *ios102* mutant seedlings with the *35S:apoaequorin* were horizontally grown on 0.6% agar containing 1/2 MS (PhytoTech, M524), 1% sucrose, and pH5.7, for nine days. The seedlings were sprayed evenly with coelenterazine solution (NanoLight, final concentration 10 μM, 0.1% Triton X-100) and incubated for 4 h at room temperature in the dark for aequorin constitution. The whole plate of seedlings was treated with 600 mM mannitol in the dark, and then imaged the $Ca^{2+}$ luminescence immediately by using a cooled charge-coupled device (Princeton Instruments, New Jersey, USA) controlled by WinView/32 (Roper).

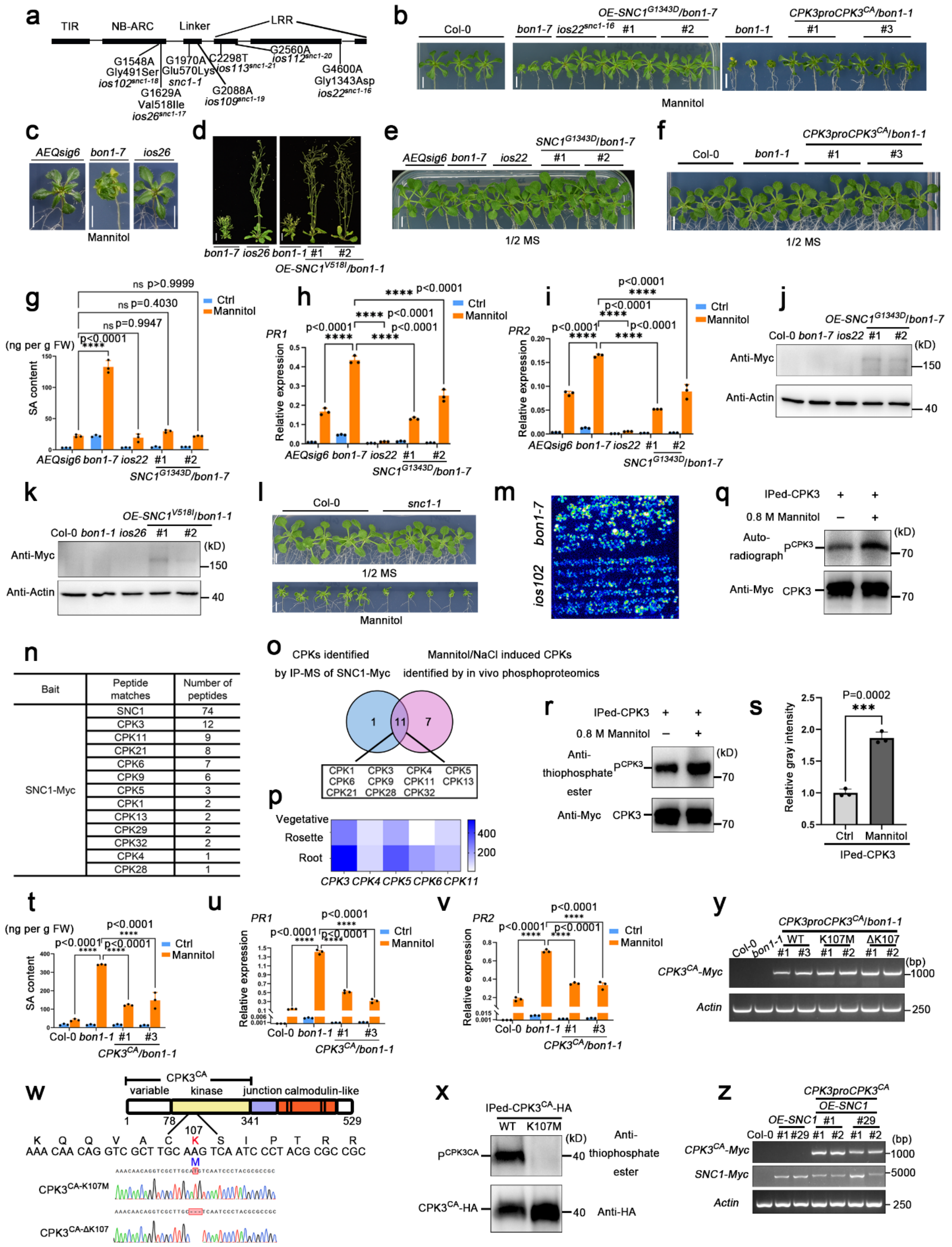
a
TIR
NB-ARC
Linker
LRR
G1548A Gly491Ser ios102snc1-18
G1629A Val518Ile ios26snc1-17
G1970A Glu570Lys snc1-1
G2088A ios109snc1-19
C2298T ios113snc1-21
G2560A ios112snc1-20
G4600A Gly1343Asp ios22snc1-16
b
Col-0
bon1-7
ios22snc1-16
OE-SNC1G1343D/bon1-7
#1
#2
bon1-1
CPK3proCPK3CA/bon1-1
#1
#3
Mannitol
c
AEQsig6
bon1-7
ios26
Mannitol
d
bon1-7
ios26
bon1-1
#1
#2
OE-SNC1V518I/bon1-1
e
AEQsig6
bon1-7
ios22
SNC1G1343D/bon1-7
#1
#2
1/2 MS
f
Col-0
bon1-1
CPK3proCPK3CA/bon1-1
#1
#3
1/2 MS
g
SA content (ng per g FW)
ns p>0.9999
ns p=0.4030
ns p=0.9947
p<0.0001 ****
Ctrl
Mannitol
AEQsig6 bon1-7 ios22 #1 #2
SNC1G1343D/bon1-7
h
PR1
Relative expression
p<0.0001 ****
Ctrl
Mannitol
AEQsig6 bon1-7 ios22 #1 #2
SNC1G1343D/bon1-7
i
PR2
Relative expression
p<0.0001 ****
Ctrl
Mannitol
AEQsig6 bon1-7 ios22 #1 #2
SNC1G1343D/bon1-7
j
OE-SNC1G1343D/bon1-7
Col-0 bon1-7 ios22 #1 #2
Anti-Myc
Anti-Actin
(kD)
150
40
k
OE-SNC1V518I/bon1-1
Col-0 bon1-1 ios26 #1 #2
Anti-Myc
Anti-Actin
(kD)
150
40
l
Col-0
snc1-1
1/2 MS
Mannitol
m
bon1-7
ios102
q
IPed-CPK3 + +
0.8 M Mannitol − +
Auto-radiograph
pCPK3
Anti-Myc
CPK3
(kD)
70
70
n
Bait | Peptide matches | Number of peptides
SNC1-Myc | SNC1 | 74
CPK3 | 12
CPK11 | 9
CPK21 | 8
CPK6 | 7
CPK9 | 6
CPK5 | 3
CPK1 | 2
CPK13 | 2
CPK29 | 2
CPK32 | 2
CPK4 | 1
CPK28 | 1
o
CPKs identified by IP-MS of SNC1-Myc
Mannitol/NaCl induced CPKs identified by in vivo phosphoproteomics
1
11
7
CPK1 CPK3 CPK4 CPK5
CPK6 CPK9 CPK11 CPK13
CPK21 CPK28 CPK32
p
Vegetative
Rosette
Root
CPK3 CPK4 CPK5 CPK6 CPK11
400
200
r
IPed-CPK3 + +
0.8 M Mannitol − +
Anti-thiophosphate ester
pCPK3
Anti-Myc
CPK3
(kD)
70
70
s
Relative gray intensity
P=0.0002
***
Ctrl
Mannitol
IPed-CPK3
t
SA content (ng per g FW)
p<0.0001 ****
Ctrl
Mannitol
Col-0 bon1-1 #1 #3
CPK3CA/bon1-1
u
PR1
Relative expression
p<0.0001 ****
Ctrl
Mannitol
Col-0 bon1-1 #1 #3
CPK3CA/bon1-1
v
PR2
Relative expression
p<0.0001 ****
Ctrl
Mannitol
Col-0 bon1-1 #1 #3
CPK3CA/bon1-1
y
CPK3proCPK3CA/bon1-1
Col-0
bon1-1
WT
K107M
ΔK107
#1 #3 #1 #2 #1 #2
CPK3CA-Myc
Actin
(bp)
1000
250
w
CPK3CA
variable
kinase
junction
calmodulin-like
1
78
341
529
107
K Q Q V A C K S I P T R R
AAA CAA CAG GTC GCT TGC AAG TCA ATC CCT ACG CGC CGC
M
CPK3CA-K107M
CPK3CA-ΔK107
x
IPed-CPK3CA-HA
WT
K107M
pCPK3CA
CPK3CA-HA
(kD)
40
40
Anti-thiophosphate ester
Anti-HA
z
CPK3proCPK3CA
OE-SNC1
OE-SNC1
Col-0 #1 #29 #1 #2 #1 #2
#1
#29
CPK3CA-Myc
SNC1-Myc
Actin
(bp)
1000
5000
250

**Extended Data Fig. 1. Dominant-negative *SNC1* mutations and constitutively active form of *CPK3* suppress NLR signaling to maintain plant growth in *bon1* mutants.**

**a**, Gene structure of *SNC1*. The mutations of *SNC1* in the *insensitive to osmotic stress* (*ios*) mutants are indicated. The *ios* mutants are suppressors of the *bon1-7* mutant based on plant growth under osmotic stress. The *ios26* (*snc1-17*, a single amino acid change V518I) and *ios102* (*snc1-18*, a single amino acid change G491S) mutations were distributed on the NB-ARC domain, probably affecting SNC1 oligomerization. The *ios22* (*snc1-16*, a single amino acid change G1343D), *ios112* (*snc1-20*, a nonsense mutation resulting in a premature termination at codon 734), and *ios113* (*snc1-21*, a single amino acid change L647F) mutations were distributed on the LRR domain, probably affecting SNC1 autoinhibition. Interestingly, the *ios109* (*snc1-19*, a single amino acid change S609N) mutation was distributed in the linker between the two domains (NL linker), adjacent to the autoimmune mutation of the gain-of-function *snc1-1* (a single amino acid change E561K). **b**,**c**,**e**,**f**, Shoot growth of Col-0, *bon1*, *ios22* (*snc1-16*) and *ios26* (*snc1-17*) mutants, and *bon1* transformed with *OE-SNC1*$^{G1343D}$ (corresponding to *snc1-16* mutation) or *CPK3pro::CPK3*$^{CA}$ 15 days after the seedlings were transferred to 1/2 MS medium (**e** and **f**) or 1/2 MS medium containing 100 mM mannitol (**b**,**c**). **d**, Morphology of six-week-old soil-grown *bon1* and *ios26* (*snc1-17*), and *OE-SNC1*$^{V518I}$ (corresponding to *snc1-17* mutation) transgenic plants in *bon1-1* background. **g**-**i**, Levels of free SA (**g**) and the relative expression of *PR1* (**h**) and *PR2* (**i**) in *AEQsig6*, *bon1-7* and *ios22*, and *bon1-7* transformed with *OE-SNC1*$^{G1343D}$ five days after the seedlings were transferred to 1/2 MS medium containing 0 or 100 mM mannitol. **j**,**k**, Protein abundance of SNC1$^{G1343D}$ (**j**, snc1-16) and SNC1$^{V518I}$ (**k**, snc1-17) in *OE- SNC1*$^{G1343D}$ and *OE-SNC1*$^{V518I}$ transgenic plants in *bon1* background was detected by western blot. The anti-Myc antibody was used to detect SNC1$^{G1343D}$ (**j**) and SNC1$^{V518I}$ (**k**). Actin was used as a loading control. **l**, Shoot growth of Col-0 (wild-type) and *snc1-1* mutant 15 days after the seedlings were transferred to the 1/2 MS medium containing 0 or 100 mM mannitol. **m**, Aequorin-based monitoring of $Ca^{2+}$ signals in *bon1-7* and *ios102* (*snc1-18*) treated with 600 mM mannitol. **n**, CPKs were co-immunoprecipitated with full-length SNC1 in *OE-SNC1-Myc* transgenic plants in Col-0 background. SNC1-Myc protein was immunoprecipitated from *OE-SNC1-Myc* transgenic seedlings treated with 600 mM mannitol. The immunoprecipitation-mass spectrometry was conducted one time as a screen. **o**, Venn diagram represents the overlap of CPKs detected as candidate SNC1 interacting proteins (**n**) and candidate osmotic stress-responsive proteins according to the *in vivo* phosphoproteomics data from (https://www.pnas.org/doi/10.1073/pnas.1308974110). **p**, Expression of *CPK3*, *4*, *5*, *6*, and *11* in

*Arabidopsis* according to the *Arabidopsis* electronic fluorescent pictograph (eFP) browser. **q**-**s**, Activation of CPK3 by hyperosmotic stress in seedlings. Myc-tagged CPK3 was immunoprecipitated from 7-day-old *CPK3pro::CPK3-4Myc* transgenic seedlings treated with 0 or 0.6 M mannitol for 30 min. The immunoprecipitated CPK3 was used for the phosphorylation assay with $^{32}$P-labeled ATP (**q**) or the ATP analog ATPγS (**r**). Autoradiography exhibits phosphorylation of CPK3 protein in (**q**). *In vitro* kinase assays were also performed using the ATP analog ATPγS. After the PNBM alkylation reaction, the thiophosphate ester groups on the substrate were detected by the anti-thiophosphate ester antibody (**r**). Total CPK3 was detected by an anti-Myc antibody. The relative gray intensity of the bands detected by anti-thiophosphate ester antibody was quantified from three independent results (**s**), representing the phosphorylation level of CPK3 in *CPK3pro::CPK3-4Myc* transgenic seedlings with or without mannitol treatment. **t**-**v**, Levels of free SA (**t**) and the relative expression of *PR1* (**u**) and *PR2* (**v**) in Col-0, *bon1-1*, and *bon1-1* transformed with *CPK3pro::CPK3*$^{CA}$ 5 days after the seedlings were transferred to 1/2 MS medium containing 0 or 100 mM mannitol. **w**, The structure of CPK3 (top) and Sanger sequencing chromatograms (bottom) indicate wild-type *CPK3* and mutations on *CPK3* in *CPK3pro::CPK3*$^{CA}$ transgenic plants in a *bon1-1* mutant background, with the kinase-dead mutations, including K107M and ΔK107. **x**, The kinase activity of CPK3-CA and kinase-dead mutation of CPK3-CA (K107M) immunoprecipitated *in planta*. HA-tagged CPK3$^{CA}$ and CPK3$^{CA\text{-}K107M}$ were immunoprecipitated from *N. benthamiana* leaves at 48 hours after transiently expressing *CPK3*$^{CA}$*-HA* or *CPK3*$^{CAK107M}$*-HA*. An anti-thiophosphate ester antibody was used to detect the autophosphorylation of CPK3$^{CA}$ or CPK3$^{CA\text{-}K107M}$ protein kinases. Total CPK3$^{CA}$ and CPK3$^{CA\text{-}K107M}$ proteins were detected by an anti-HA antibody. **y**, Expression of *CPK3*$^{CA}$*-Myc*, *CPK3*$^{CA\text{-}K107}$*-Myc*, *CPK3*$^{CA\text{-}\Delta K107}$*-Myc* and *Actin* in Col-0, *bon1-1* mutant, and transgenic plants harboring *CPK3pro::CPK3*$^{CA}$, *CPK3pro::CPK3*$^{CA\text{-}K107M}$, and *CPK3pro::CPK3*$^{CA\text{-}\Delta K107}$ in *bon1-1* background. **z**, Expression of *SNC1-Myc*, *CPK3*$^{CA}$*-Myc*, and *Actin* in Col-0, and *OE-SNC1-4Myc* transgenic plants with or without *CPK3pro::CPK3*$^{CA}$ in Col-0 background. Scale bars, 0.5 cm in (**b**,**c**,**e**,**f**,), and 2 cm in (**d**). Error bars represent means ± SD (n = 3 experiments in **g**-**i** and **t**-**v**). The statistical analysis was performed using a two-way ANOVA with Tukey's test. All experiments except (**n**, one time; **o** and **p**, previously published data) were repeated at least three times with similar results.

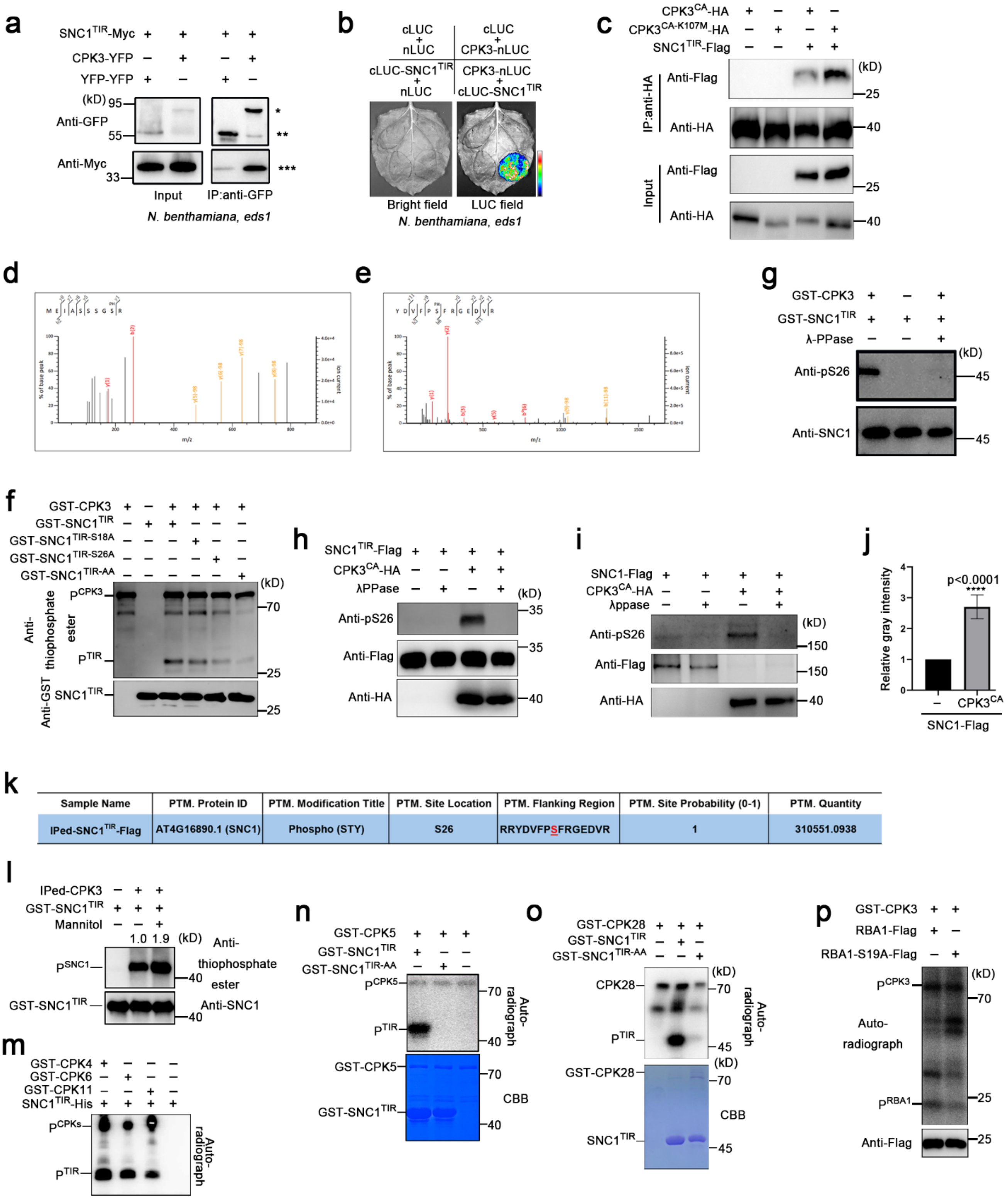


**Fig. S2. CPKs interact with and phosphorylate SNC1$^{TIR}$ both in *vitro* and in *planta*.**

**a**, CPK3-SNC1$^{TIR}$ interaction in *N. benthamiana eds1* mutant leaves in a co-immunoprecipitation assay. *SNC1$^{TIR}$-Myc*, *CPK3-YFP*, and double *YFP* were transiently expressed in *N. benthamiana eds1* mutant leaves. Total proteins were extracted and immunoprecipitated with anti-GFP agarose and detected with an anti-Myc antibody. *, CPK3-YFP; **, YFP-YFP; ***, SNC1$^{TIR}$-Myc. **b**, CPK3-SNC1$^{TIR}$ interaction in *N. benthamiana eds1* mutant leaves using split-LUC assays. CPK3 and SNC1$^{TIR}$ were fused to the split N- or C-terminal fragments of LUC (CPK3-nLUC and cLUC- SNC1$^{TIR}$). Unfused nLUC and cLUC were used as negative controls. **c**, CPK3$^{CA}$-SNC1$^{TIR}$ or CPK3$^{CA-K107M}$-SNC1$^{TIR}$ interaction in *N. benthamiana* leaves in a co-immunoprecipitation assay. *SNC1$^{TIR}$-Flag*, *CPK3$^{CA}$-HA* and *CPK3$^{CA-K107M}$-HA* were transiently expressed in *N. benthamiana* leaves. Total proteins were extracted and immunoprecipitated with anti-HA beads and detected with an anti-Flag antibody. **d**,**e**, Mass spectrometric analysis of SNC1 phosphorylation at Ser18 (**d**) and Ser26 (**e**) by CPK3. The purified recombinant GST-SNC1$^{TIR}$ protein was phosphorylated by the recombinant GST-CPK3 protein kinase and the phosphosites were detected by mass spectrometric analysis. **f**, Phosphorylation of wild-type and mutated recombinant SNC1$^{TIR}$ by recombinant CPK3. An anti-thiophosphate ester antibody was used to detect the phosphorylation of wild-type and mutated SNC1$^{TIR}$ and CPK3. The anti-GST antibody was used to detect the loading of GST-SNC1$^{TIR}$, GST-SNC1$^{TIR-S18A}$, GST-SNC1$^{TIR-S26A}$, and GST-SNC1$^{TIR-AA\ (S18A/S26A)}$. **g**, CPK3 phosphorylates SNC1$^{TIR}$ at Ser26. *In vitro* kinase assays were performed by incubating GST-SNC1$^{TIR}$ with GST-CPK3 at kinase buffer without $^{32}$P-labeled ATP, without or with λ-protein phosphatase (λ-PPase). Anti-phospho-Ser26-SNC1$^{TIR}$ antibodies were used to detect the phosphorylation of SNC1$^{TIR}$. An anti-SNC1 antibody was used to detect total recombinant GST-SNC1$^{TIR}$. **h**-**j**, Phosphorylation of Ser26 of SNC1$^{TIR}$-Flag (**h**) and full-length SNC1-Flag (**i**) by CPK3$^{CA}$-HA *in planta*. Flag-tagged SNC1$^{TIR}$-Flag or full-length SNC1-Flag was immunoprecipitated from *N. benthamiana* leaves co-expressed *SNC1$^{TIR}$-Flag* or full-length *SNC1-Flag* and *CPK3$^{CA}$-HA*. Anti-phospho-Ser26-SNC1$^{TIR}$ antibody (anti-pS26) was used to detect the phosphorylation of SNC1$^{TIR}$ (**h**) and full-length SNC1 (**i**). Anti-Flag and anti-HA antibodies were used as the loading control. The relative gray intensity of the bands detected by anti-pS26 was quantified from three independent results (**j**). **k**, SNC1$^{TIR}$ phosphorylation at Ser26 by CPK3$^{CA}$ *in planta* was detected by the data-independent acquisition (DIA) based mass spectrometry analysis. Flag-tagged SNC1$^{TIR}$ was immunoprecipitated from *N. benthamiana* leaves co-expressed *SNC1$^{TIR}$-Flag* and *CPK3$^{CA}$-HA*. The immunoprecipitated SNC1$^{TIR}$ was used for the DIA-based mass spectrometry assay. **l**, Phosphorylation of SNC1$^{TIR}$ by CPK3 immunoprecipitated from seedlings treated with hyperosmotic stress. Myc-tagged CPK3 was immunoprecipitated from 7-day-old *CPK3pro::CPK3-4Myc* transgenic seedlings treated with 0.6 M mannitol for 30 min and used for the phosphorylation assay. An anti-thiophosphate ester antibody was used to detect the phosphorylated SNC1$^{TIR}$. An anti-SNC1 antibody was used to detect total SNC1$^{TIR}$. **m**, Phosphorylation of SNC1$^{TIR}$ by recombinant GST-CPK4, GST-CPK6, and GST-CPK11. Autoradiography exhibits phosphorylation of GST-CPK4, GST-CPK6, GST-CPK11, and SNC1$^{TIR}$-His proteins. **n**, Phosphorylation of SNC1$^{TIR}$ and SNC1$^{TIR\text{-}S18A/S26A\ (AA)}$ by recombinant GST-CPK5. Autoradiography (top) and Coomassie staining (bottom) exhibit phosphorylation and the loading of recombinant GST-CPK5, GST-SNC1$^{TIR}$, and GST-SNC1$^{TIR\text{-}S18AS26A\ (AA)}$. **o**, Phosphorylation of SNC1$^{TIR}$ and SNC1$^{TIR\text{-}S18A/S26A\ (AA)}$ by recombinant GST-CPK28. Autoradiography (top) and Coomassie staining (bottom) exhibit phosphorylation and the loading of recombinant GST-CPK28, GST-SNC1$^{TIR}$, and GST-SNC1$^{TIR\text{-}S18AS26A\ (AA)}$. **p**, Phosphorylation of RBA1 and RBA1S19A by recombinant GST-CPK3. Autoradiography (top) exhibits phosphorylation of RBA1-Flag, RBA1S19A-Flag, and recombinant GST-CPK3. The anti-Flag antibody was used as a loading control (bottom). All experiments except (**d**, **e**, **k**, one time) were repeated at least three times with similar results.

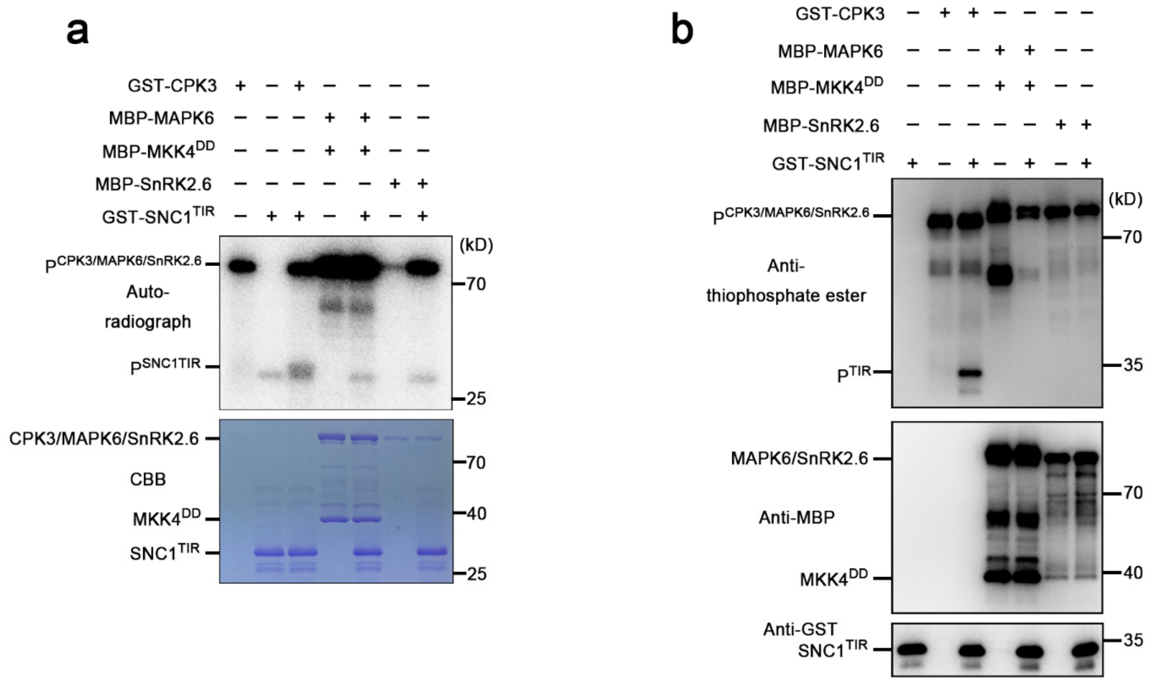


**Fig. S3. MPK6 and SnRK2.6 cannot phosphorylate the TIR domain of SNC1.**

**a**,**b**, Phosphorylation of wild-type SNC1$^{TIR}$ by CPK3, MPK6 and SnRK2.6. GST-CPK3 was used as a positive control. Autoradiography (top) and Coomassie staining (bottom) exhibit phosphorylation and the loading of recombinant MBP-MAPK6, MKK4$^{DD}$, GST-CPK3, MBP-SnRK2.6 and GST-SNC1$^{TIR}$ (**a**). An anti-thiophosphate ester antibody was used to detect the phosphorylation of SNC1$^{TIR}$, CPK3, MAPK6 and SnRK2.6 (**b**, top). The anti-MBP and anti-GST antibodies were used to detect the loading of recombinant GST-SNC1$^{TIR}$, MBP-MAPK6, MBP-MKK4$^{DD}$, and MBP-SnRK2.6 (**b**, middle and bottom). All experiments were repeated at least three times with similar results.

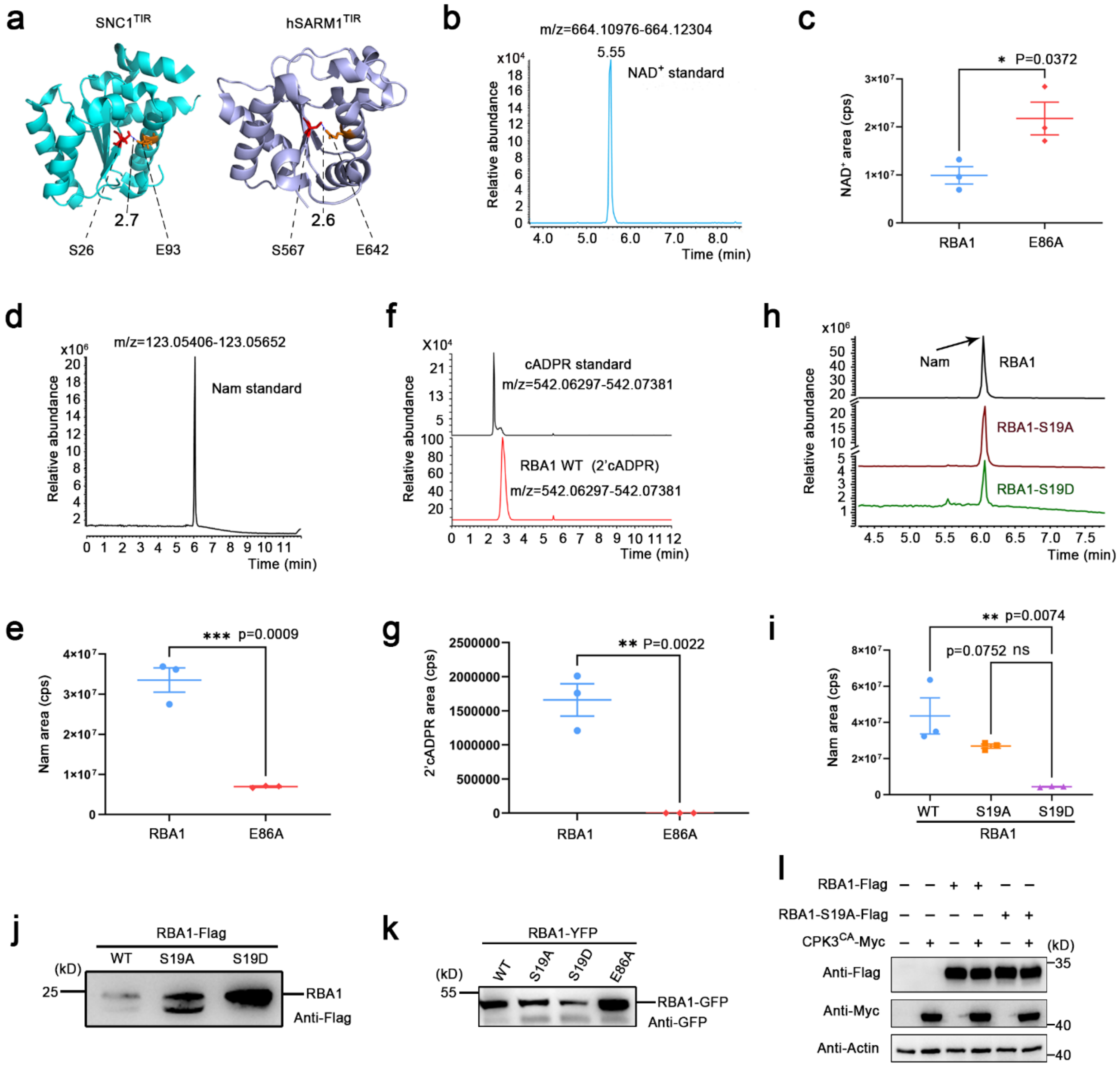


**Fig. S4. Phosphorylation of TIR represses its NADase activity.**

**a**, Cartoon representation of the SNC1$^{TIR}$ (PDB: 5TEC) and human SARM1$^{TIR}$ (PDB: 8D0J) structures highlighting the conserved serines and catalytic glutamates. The catalytic glutamate (orange) and the conserved serine (red) in the catalytic pockets are presented as sticks. The distances are shown for the carboxylic acid group of the catalytic glutamate with the hydroxyl group of the conserved serine. **b**-**g**, Purified wild-type RBA1 protein, but not the catalytically dead RBA1-E86A mutant, exhibited strong NADase enzyme activity. LC-MS/MS results showing that RBA1 could degrade NAD$^+$ (**b** and **c**) and produce Nam (**d** and **e**) and 2′cADPR (**f** and **g**). LC-MS/MS chromatograms (**b**, **d**, and **f**) and areas under curves (**c**, **e**, and **g**) indicating that the NADase enzyme activity of RBA1 depends on the catalytic glutamate. The purified RBA1-S19A and RBA1-S19D proteins were also used for *in vitro* NADase enzyme assays (Fig. 2, c-f). **h**,**i**, LC-MS/MS showing the generation of Nam by wild-type RBA1, RBA1-S19A, and RBA1-S19D *in vitro*. LC-MS/MS chromatograms (**h**) and areas under curves (**i**) indicating that RBA1-S19D could not produce Nam. Error bars represent means ± SEM (n = 3 experiments). The statistical analysis was performed using a two-tailed Student's t-test in (**c**), (**e**), and (**g**), or a one-way ANOVA with Tukey's test in (**i**). LC-MS/MS chromatograms and areas under curves also indicating that RBA1-S19D could not degrade $NAD^+$ (Fig. 2, c and d) and produce 2′cADPR (Fig. 2, e and f). **j**, Wild-type and mutated RBA1-Flag proteins were detected by an anti-Flag antibody. Recombinant RBA1 proteins were expressed in insect cells and purified using anti-Flag beads. The purified RBA1 proteins were used for *in vitro* NADase enzyme assays (Fig. 2, c-f, and fig. S4, b-i). **k**, Wild-type and mutated RBA1-YFP proteins were expressed in *N. benthamiana eds1-2* mutant and detected by an anti-GFP antibody. The resulting accumulation of 2′cADPR was shown in Figure 2g. **l**, *RBA1-Flag* and *RBA1-S19A-Flag* were stably expressed in the HEK293T cells, with transient expression of *$CPK3^{CA}$-My*c. The protein abundance of RBA1-Flag, RBA1-S19A-Flag and $CPK3^{CA}$-Myc was detected by western blot, using the anti-Flag and anti-Myc antibodies. Actin was used as a loading control, and it was detected with the anti-actin antibody. The resulting degradation of $NAD^+$ was shown in Figure 2h. All experiments except (**a**) were repeated at least three times with similar results.

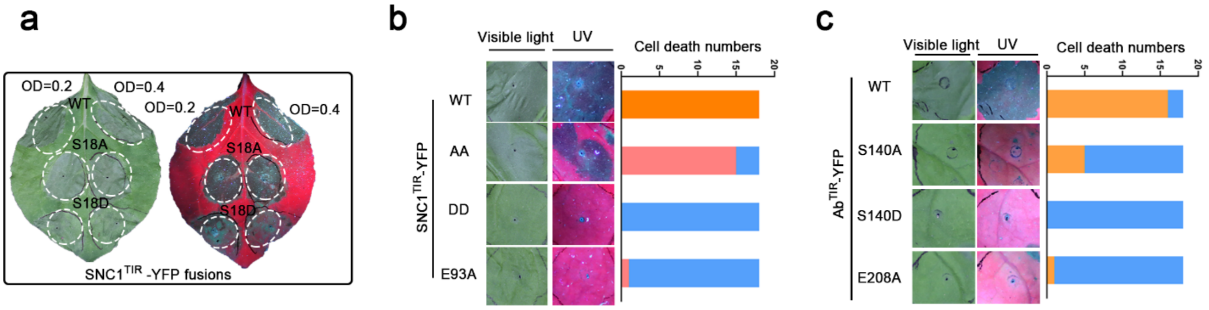


**Fig. S5. Phosphomimetic mutation of TIR represses TIR-mediated cell death.**

Cell death phenotypes of wild-type and mutated TIRs upon transient expression in *N. benthamiana* leaves. The cell-death phenotype was visualized under visible light (left panel) or UV (right panel) and imaged three days after agroinfiltration inoculation. Different degrees of cell death were defined as the cell death score (0, blue; 1, coral pink; 2, orange). Each color on stacked bars shows the proportions (in numbers) of the corresponding degrees of cell death. **a**, Expression of $SNC1^{TIR}$-*YFP*, non-phosphorylatable $SNC1^{TIR\text{-}S18A}$-*YFP*, and phospho-mimic $SNC1^{TIR\text{-}S18D}$-*YFP*. Plant leaves were infiltrated with *Agrobacterium tumefaciens* bacteria with OD600 at 0.2 (left) or 0.4 (right). **b**, Expression of $SNC1^{TIR}$-*YFP*, non-phosphorylatable $SNC1^{TIR\text{-}AA\ (S18A/S26A)}$-*YFP*, phospho-mimic $SNC1^{TIR\text{-}DD\ (S18D/S26D)}$-*YFP*, and catalytically dead $SNC1^{TIR\text{-}E84A}$-*YFP*. **c**, Expression of $Ab^{TIR}$-*YFP*, non-phosphorylatable $Ab^{TIR\text{-}S140A}$-*YFP*, phospho-mimic $Ab^{TIR\text{-}S140D}$-*YFP*, and catalytically dead $Ab^{TIR\text{-}E208A}$-*YFP*. All experiments were repeated at least three times with similar results.

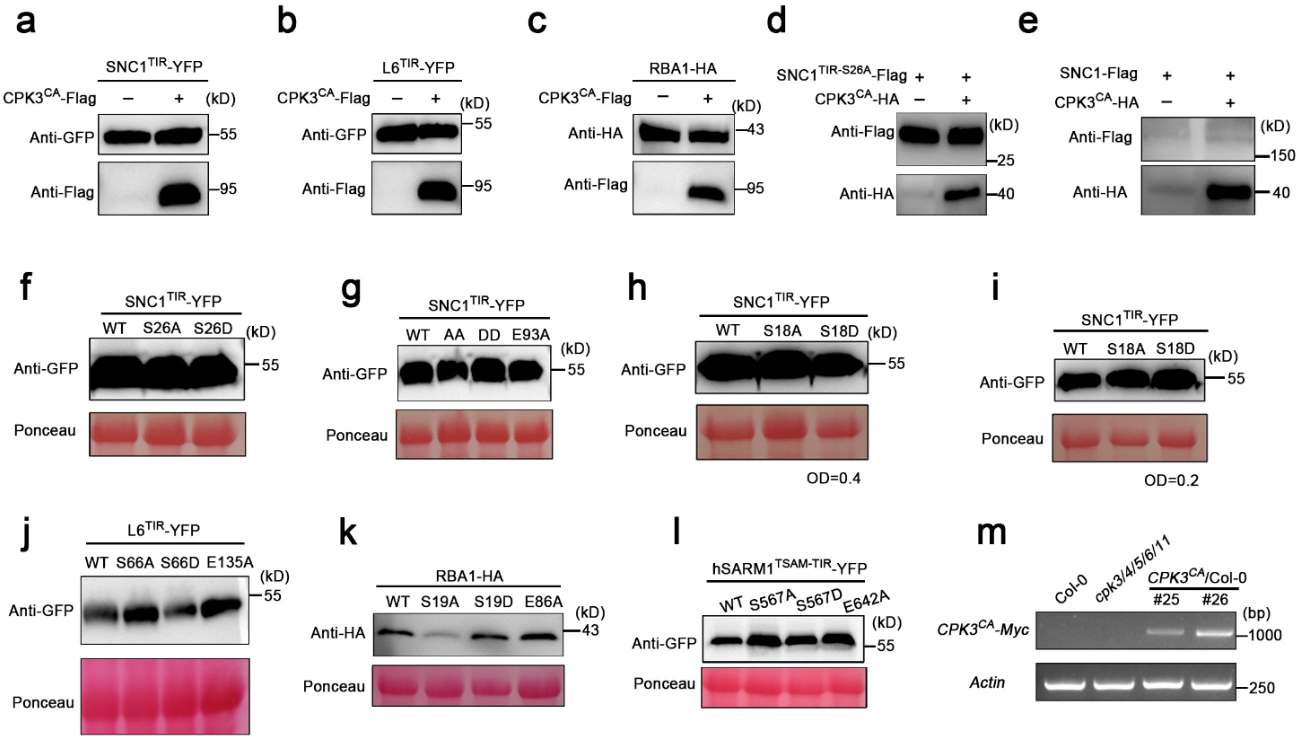


**Fig. S6. Expression of wild-type and mutated TIR proteins and CPKs *in planta*.**

**a**-**c**, SNC1$^{TIR}$-YFP, L6$^{TIR}$-YFP, RBA1-HA, and CPK3$^{CA}$-cLUC-Flag proteins were transiently expressed in *N. benthamiana* leaves and detected by western blot. The anti-GFP antibody was used to detect SNC1$^{TIR}$-YFP and L6$^{TIR}$-YFP. The anti-HA antibody was used to detect RBA1-HA. The anti-Flag antibody was used to detect CPK3$^{CA}$-cLUC-Flag. Cell death phenotypes of wild-type TIRs with or without CPK3$^{CA}$ were shown in Figure 3b. **d**,**e**, SNC1$^{TIR\text{-}S26A}$-Flag, full-length SNC1-Flag and CPK3$^{CA}$-HA were transiently expressed in *N. benthamiana* leaves and detected by western blot. The anti-Flag antibody was used to detect SNC1$^{TIR}$-Flag or full-length SNC1-Flag. The anti-HA antibody was used to detect CPK3$^{CA}$-HA. Cell death phenotypes of SNC1$^{TIR\text{-}S26A}$ and full-length SNC1 with or without CPK3$^{CA}$ were shown in Figures 3c and 3d, respectively. **f**-**l**, Wild-type and mutated TIR proteins were transiently expressed in *N. benthamiana* leaves and detected by western blot. Total proteins were extracted from *N. benthamiana* leaves. Staining of RuBisCO with Ponceau S was used as a loading control. Anti-GFP antibody was used to detect wild-type and mutated TIR-YFP proteins (**f**-**j**, and **l**). Anti-HA antibody was used to detect RBA1-HA (**k**). SNC1$^{TIR}$ (**f**-**i**) related cell death phenotype was shown in Figures 3e and S5; L6$^{TIR}$ (**j**) related cell death phenotype was shown in Figure 3f; RBA1 (**k**) related cell death phenotype was shown in Figure 3g; while hSARM1$^{tSAM\text{ -}TIR}$ (**l**) related cell death phenotype was shown in Figure 3h. **m**, Expression of *CPK3*$^{CA}$*-Myc* and *Actin* in Col-0, *cpk3/4/5/6/11* quintuple mutant, and *CPK3pro::CPK3*$^{CA}$ transgenic plants in Col-0 background using reverse transcriptional PCR analysis. The corresponding plant growth phenotype was shown (Fig. 3, l and m). All experiments were repeated at least three times with similar results.

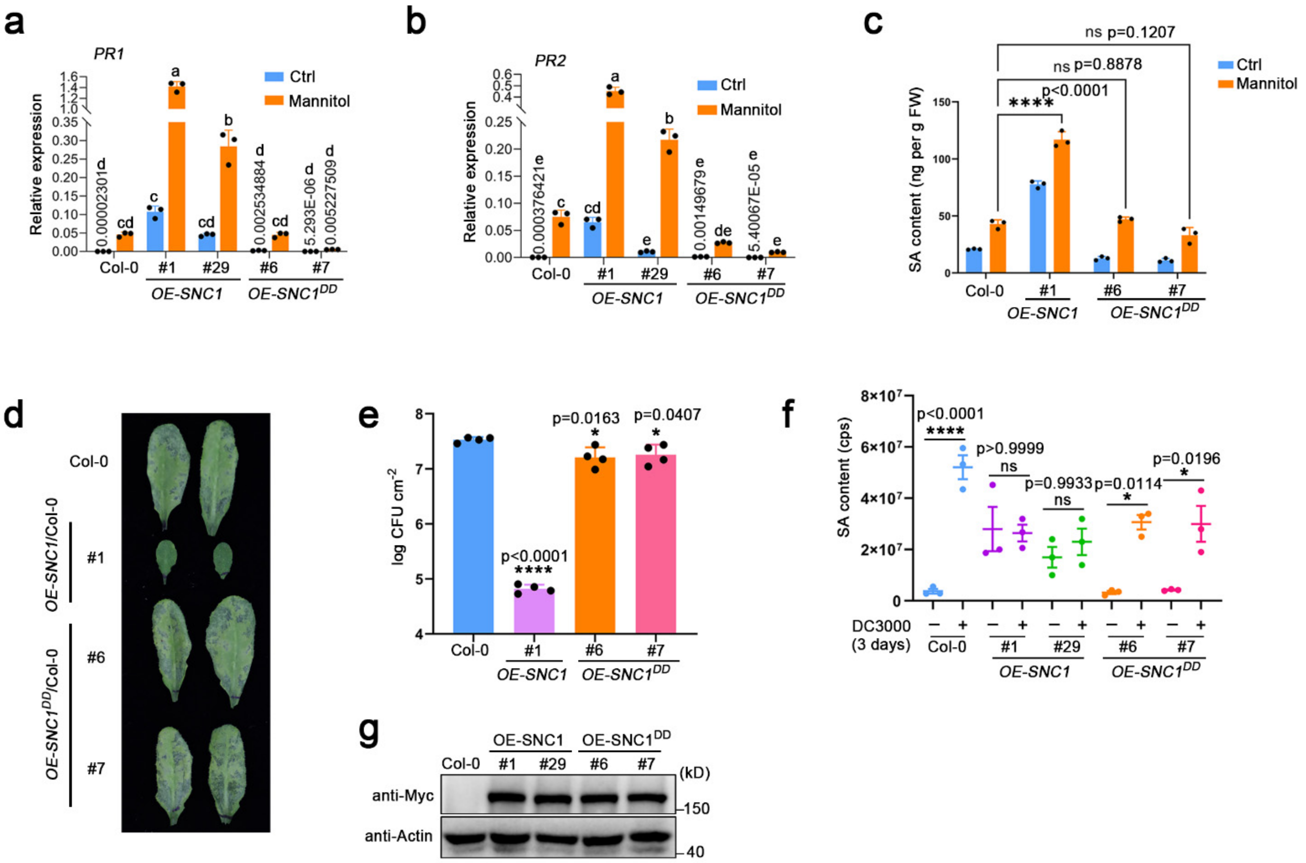


**Fig. S7. Phosphomimetic mutation of SNC1$^{TIR}$ represses its immune function and maintains plant growth.**

**a**-**c**, Levels of relative expression of *PR1* (**a**), *PR2* (**b**) and free SA (**c**) in Col-0 and *OE-SNC1-Myc* transgenic plants expressing wild-type or *S18D/S26D* (*DD*)-mutated *SNC1* in Col-0 background five days on the 1/2 MS medium containing 0 or 100 mM mannitol. **d-f**, *Pst* DC3000 triggered plant immunity in Col-0 and *OE-SNC1-4Myc* transgenic plants expressing wild-type or *S18D/S26D* (*DD*)-mutated *SNC1* in Col-0 background. Plant leaves were infiltrated with *Pst* DC3000 bacteria with OD600 at 0.002. Disease symptoms (**d**), bacterial populations (**e**), and levels of free SA (**f**) were monitored 3 days post-infection. Values are means ± SD (n = 3 experiments) in (**a** and **b**), with different letters indicating significant differences as evaluated by post-hoc Tukey tests after two-way ANOVAs ($p < 0.05$). Error bars represent means ± SD (n = 3 experiments) in (**c**), ± SD (n = 4 experiments) in (**e**), and ± SEM (n = 3 experiments) in (**f**). The statistical analysis was performed using a two-way ANOVA with Tukey's test in (**c** and **f**), and a one-way ANOVA with Tukey's test in (**e**). **g**, Protein abundance of SNC1 and SNC1$^{S18D/S26D\ (DD)}$ in *OE-SNC1-4Myc* transgenic plants expressing wild-type or *S18D/S26D*

(*DD*)-mutated *SNC1* in Col-0 background. The anti-Myc antibody was used to detect SNC1 and SNC1$^{S18DS26D\ (DD)}$. All experiments were repeated at least three times with similar results.

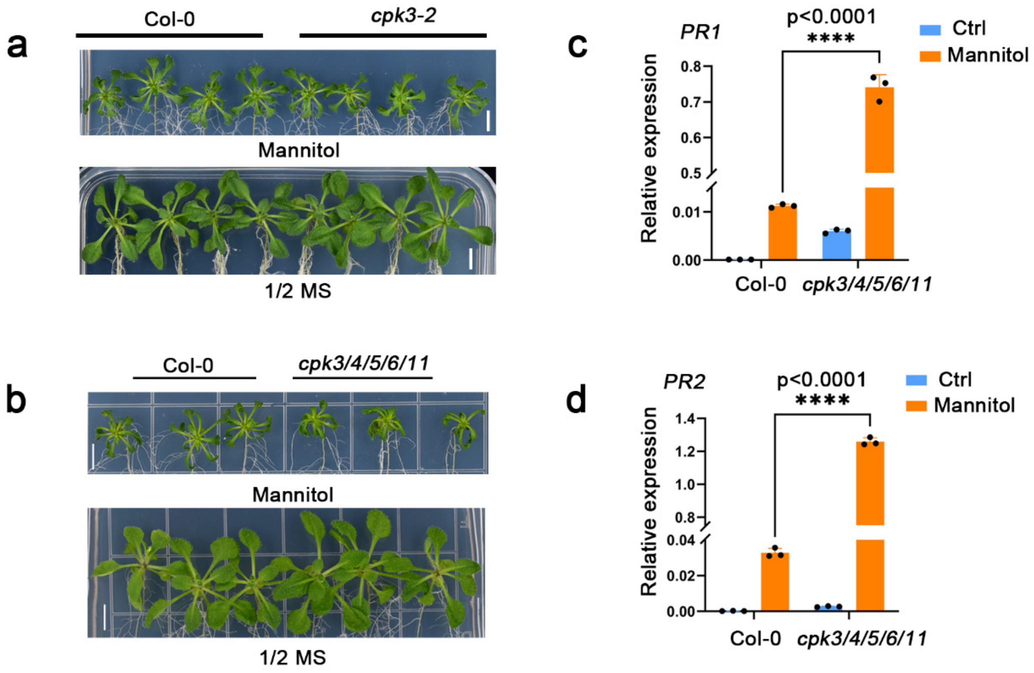


**Fig. S8. The *cpk3/4/5/6/11* quintuple mutant exhibited shoot growth defects and elevated levels of defense marker gene expression under hyperosmotic stress.**

**a**,**b**, Shoot growth of Col-0, *cpk3-2* mutant (**a**), and *cpk3/4/5/6/11* quintuple mutant (**b**), 15 days after the seedlings were transferred to the 1/2 MS medium containing 0 mM or 100 mM mannitol. Scale bars, 0.5 cm. **c**,**d**, Levels of relative expression of *PR1* (**c**) and *PR2* (**d**) in Col-0 and *cpk3/4/5/6/11* quintuple mutant five days on the 1/2 MS medium containing 0 or 100 mM mannitol. All experiments were repeated at least three times with similar results.

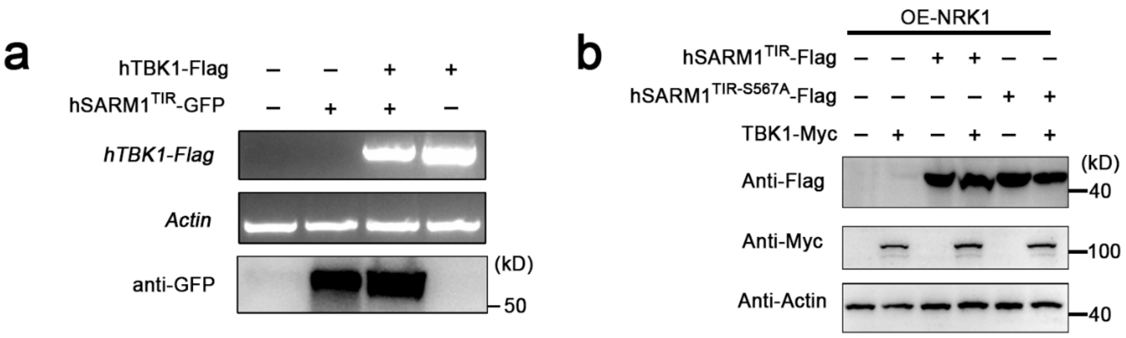


**Fig. S9. Regulation of SARM1$^{TIR}$ by TBK1.**

**a**, Expression of *hTBK1-Flag* and *Actin* in *N. benthamiana* leaves transiently expressing *hSARM1$^{TIR}$-GFP* with or without *hTBK1-Flag* using reverse transcriptional PCR analysis. The anti-GFP antibody was used to detect hSARM1$^{TIR}$-GFP proteins. **b**, Human SARM1$^{TIR}$-GFP-Flag and hSARM1$^{TIR-S567A}$-GFP-Flag proteins were transiently expressed in the *NRK1*-HEK293T or *TBK1/NRK1*-HEK293T stable cell lines and detected by western blot. The anti-Flag antibody was used to detect wild-type and mutated hSARM1$^{TIR}$-GFP-Flag. The anti-Myc antibody was used to detect TBK1-Myc. The anti-actin was used as a loading control. All experiments were repeated at least three times with similar results.
